# F-actin and Myosin F control apicoplast elongation dynamics which drive apicoplast-centrosome association in *Toxoplasma gondii*

**DOI:** 10.1101/2023.01.01.521342

**Authors:** Parvathi Madhavi Devarakonda, Valeria Sarmiento, Aoife T. Heaslip

**Affiliations:** Department of Molecular and Cell Biology, University of Connecticut, Storrs, CT, 06269

**Author notes:** Address correspondence to Aoife Heaslip.

## Abstract

*Toxoplasma gondii* contains an essential plastid organelle called the apicoplast that is necessary for fatty acid, isoprenoid, and heme synthesis. Perturbations affecting apicoplast function or inheritance lead to parasite death. The apicoplast is a single copy organelle and therefore must be divided so that each daughter parasite inherits an apicoplast during cell division. In this study we identify new roles for F-actin and an unconventional myosin motor, TgMyoF, in this process. First, loss of TgMyoF and actin lead to an accumulation of apicoplast vesicles in the cytosol indicating a role for this actomyosin system in apicoplast protein trafficking or morphological integrity of the organelle. Second, live cell imaging reveals that during division the apicoplast is highly dynamic, exhibiting branched, U-shaped and linear morphologies that are dependent on TgMyoF and actin. In parasites where movement was inhibited by the depletion of TgMyoF, the apicoplast fails to associate with the parasite centrosomes. Thus, this study provides crucial new insight into mechanisms controlling apicoplast-centrosome association, a vital step in the apicoplast division cycle, which ensures that each daughter inherits a single apicoplast.

## INTRODUCTION

The phylum Apicomplexa contains over 6,000 parasites, many of which are of significant medical and veterinary importance, including *Toxoplasma gondii* the causative agent of Toxoplasmosis and *Plasmodium spp*. which causes malaria [1]. These parasites are distinguished by the presence of vestigial non-photosynthetic plastid organelle named the apicoplast (<u>Apico</u>mplexan <u>plast</u>id), that is found in most Apicomplexa.

The apicoplast is an essential organelle required for fatty acid, isoprenoid, iron-sulfur cluster and heme synthesis [2–8]. Genetic manipulations or perturbation of the apicoplast with pharmacological compounds results in apicoplast inheritance defects and parasite death [9–11]. Despite containing a small 35kb genome, the majority of apicoplast localized proteins are encoded in the nucleus. Therefore, trafficking of these proteins to the apicoplast is a critical cellular process and our understanding of the mechanisms of apicoplast protein trafficking is incomplete. Many apicoplast proteins contain a bipartite N-terminal signal sequence that includes a signal peptide (SP), which directs protein translation to the endoplasmic reticulum (ER), and an apicoplast transit peptide which is revealed after removal of the SP [12,13]. It is unclear how these proteins translocate from the ER to the apicoplast. A subset of apicoplast proteins that localize within the apicoplast membranes, including APT1, Atrx1 and FtsH1, do not contain a signal peptide or apicoplast targeting sequence and have been observed in vesicles that are thought to directly traffic these proteins from the ER to the apicoplast in a cell cycle dependent manner [14–19]. Proteins that localize to both the apicoplast and the mitochondria, are translocated via the Golgi and trafficking of these proteins to the apicoplast is perturbed by the addition of the ER retention motif HDEL [20]. Overall, there are numerous mechanisms controlling the trafficking of proteins to the apicoplast and the molecular details of each system are poorly understood.

Evolutionarily, the apicoplast was acquired by the parasite via a secondary endosymbiosis of red algae and contains four membranes [21]. The inner two membranes are derived from the progenitor chloroplast (inner most), the third from the algae plasma membrane (periplastid) and outermost membrane from plasma membrane of the Apicomplexan ancestor which first acquired this organelle (Fig. 1A). Protein translocons Tic and Toc in the inner two apicoplast membranes facilitate protein transport from the periplastid to the apicoplast lumen [22,23]. Repurposed ER-associated degradation (ERAD) proteins translocate proteins across the periplastid membrane [24,25]. How proteins traverse the outer membrane is not understood but, in some instances, may involve fusion of the outer membrane with ER derived vesicles [26].

**Figure 1.**
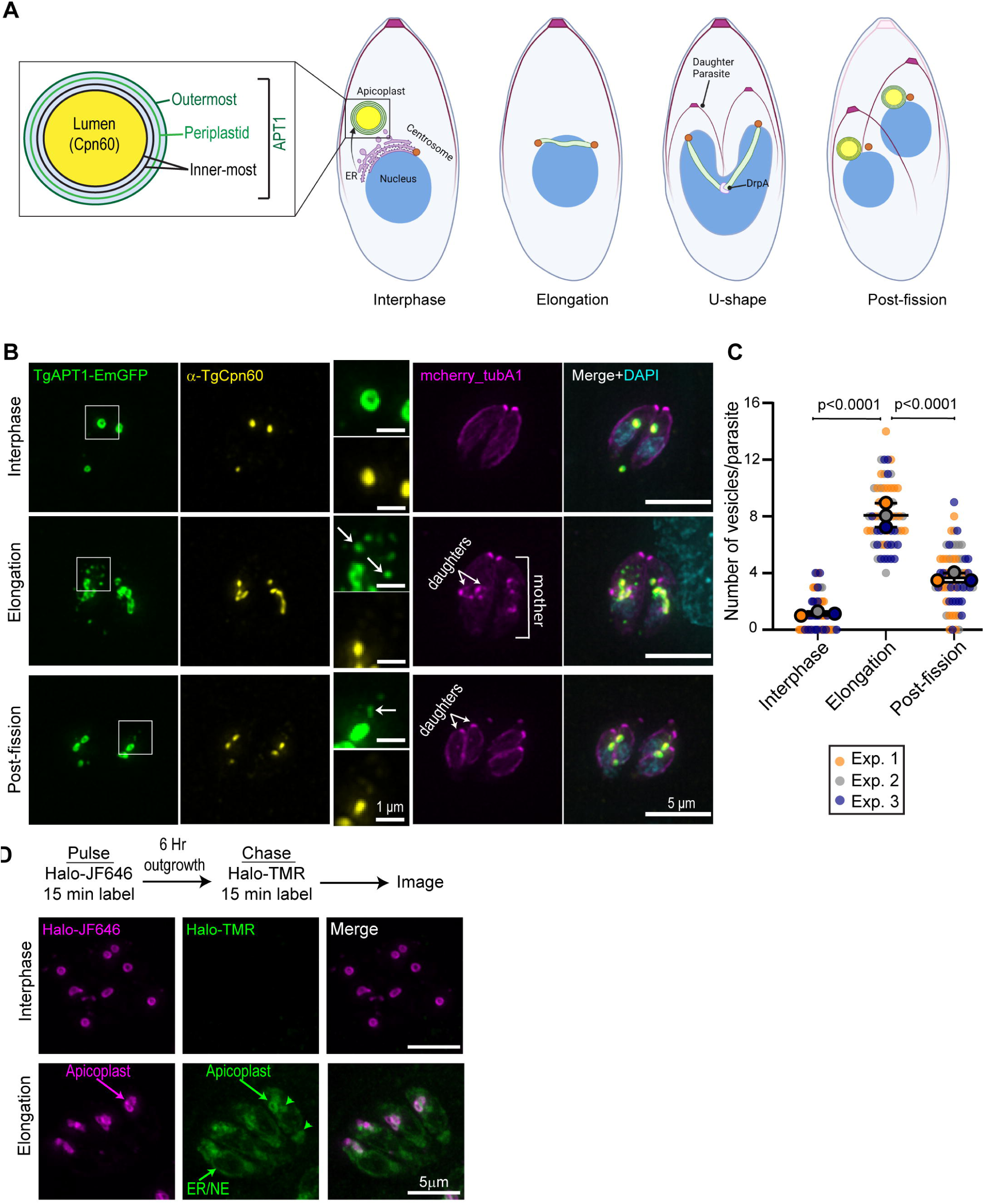
TgAPT1 vesicle trafficking occurs in the elongation phase of apicoplast division. (A) Schematic of the apicoplast division cycle in *T. gondii*. During interphase the apicoplast is circular and located at the parasite’s apical end. At the start of the cell division, apicoplast undergoes elongation and associates with a pair of duplicated centrosomes. Centrosome movement into daughter parasites, results in a U-shaped apicoplast. Membrane fission is driven by TgDrpA and results in a single apicoplast being inherited by each daughter cell. Inset: Depiction of apicoplast morphology. The apicoplast contains 4 membranes: outermost, periplastid (green) and two innermost membrane (black). TgAPT1 localizes to the apicoplast membranes. Cpn60 localizes to the apicoplast lumen. B) Maximum intensity projection of deconvolved epifluorescent images of the parasites in interphase, elongation and post-fission stages of the apicoplast division cycle. Parasites are expressing TgAPT1-EmGFP (green) and mCherry-TubulinA1 (a marker for mother and daughter parasite pellicle shown in magenta). Parasites were fixed and stained with an anti-Cpn60 antibody (a marker for the apicoplast lumen shown in yellow) and DAPI (cyan). Scale bar represents 5 μm. Inset: Brightness and contrast of insets was adjusted to facilitate visualization of dimmer TgAPT1 vesicles (white arrows). Inset scale bar represents 1 μm. (C) Quantification of the number of TgAPT1-EmGFP vesicles per parasite at interphase, elongation, and post-fission from three independent experiments (N=60 parasites). The mean from each experiment is indicated with large circles. These values were used to calculate the mean, standard error of the mean and p-value. Raw data is shown with smaller colored circles. Experiment 1 in orange, experiment 2 in grey, experiment 3 in blue. Statistical significance was determined by unpaired Student’s *t* test. (D) Pulse-chase assay to track APT1 protein synthesis. *Upper:* Assay outline. Parasites expressing APT1-Halo were labeled with JF646 for 15 minutes (pulse). After a 6 hour out-growth the parasites were labeled with Halo-TMR for 15 minutes (chase). *Lower:* Maximum intensity projections of deconvolved images of Halo-APT1 expressing parasites after Halo pulse-chase assay. No TMR labeling was observed in parasites in interphase, indicated by circular apicoplasts, demonstrating that APT1 is not expressed in interphase parasites. During apicoplast elongation, TMR-Halo staining was observed in the ER and nuclear envelope (NE) and in the apicoplast. Green arrow heads indicate accumulations of APT1-Halo/TMR in the ER at the apical and basal sides of the nucleus. Images are maximum intensity projections of deconvolved images. Scale bar represents 5 μm.

The apicoplast is a single-copy organelle, thus apicoplast division and inheritance must occur with high accuracy to ensure parasite survival. Apicoplast division is a multistep process. During interphase the apicoplast has a circular morphology and is located at the parasite’s apical end (Fig. 1A). At the beginning of cell division, the apicoplast interacts with the duplicated centrosomes and elongates [27]. This interaction between centrosomes and the apicoplast is dependent on the autophagy related protein Atg8 which localizes to the outer membrane of the apicoplast and disruption of Atg8 in both *Toxoplasma* and *Plasmodium* results in apicoplast inheritance defects [28–31]. While the apicoplast is elongated, daughter cell construction within the parasite cytosol is initiated [32]. The apical tip of the growing daughters is connected to the centrosomes via a fiber containing striated fiber assemblins (SFAs) [33], ensuring that each daughter cell inherits a single centrosome. As the centrosome migrates into the growing daughter scaffolds, the elongated apicoplast then extends to a U-shaped structure, whereupon apicoplast fission is mediated by dynamin-related protein A (DrpA) [34].

Three cytoskeletal proteins, actin (TgAct1), Myosin F (TgMyoF: an unconventional apicomplexan specific myosin motor), and the actin nucleator formin-2 also play major roles in apicoplast division [35–38]. Since TgMyoF has also been implicated in centrosome positioning, it was previously proposed that TgMyoF maintains the duplicated centrosomes in close proximity, thus allowing the apicoplast to associate with both structures [37]. However, it has also been shown that TgMyoF and actin control the transport of a variety of vesicles, including dense granules and Rab6 vesicles [38,39], thus an alternative hypothesis is that TgMyoF and actin are needed for the trafficking of vesicles to the apicoplast, which could provide new material (proteins and lipids) to the apicoplast prior to division.

In this study, we investigate the roles of TgMyoF and actin in apicoplast division. We demonstrate that vesicles containing the apicoplast protein APT1 are formed in a cell cycle dependent manner consistent with previous work [14]. Loss of MyoF and actin lead to an increased number of vesicles in the cytosol but this cytoskeletal system does not play a major role in vesicle movement. While APT1 protein translation at the ER coincides with vesicle formation, we could not definitely determine that these vesicles were ER-derived vesicles trafficking to the apicoplast.

Live cell imaging revealed that elongated apicoplasts are highly dynamic, exhibiting linear, branched and U-shaped morphologies which are driven by both TgMyoF and actin. Loss of this actomyosin system results in reduced apicoplast length and defects in apicoplast-centrosome association. Thus, we conclude that MyoF and actin play a vital role in recruiting the apicoplast to the duplicated centrosomes during apicoplast division, a vital step that ensures each daughter inherits a single apicoplast.

## MATERIAL AND METHODS

### Cell culture

RH parasites were maintained by passage in human foreskin fibroblast (HFF) cells obtained from ATCC (Catalog # CRL-2429). HFFs were grown to confluence in Dulbecco’s modified Eagle’s medium containing 10% (v/v) heat inactivated fetal bovine serum (FBS) and 1% (v/v) antibiotic/antimycotic at 37°C, 5% CO_2_. Before parasite addition, confluent HFF monolayers were maintained in Dulbecco’s modified Eagle’s medium containing 1% (v/v) heat inactivated fetal bovine serum (FBS) and 1% (v/v) antibiotic/antimycotic 37°C, 5% CO_2_.

### Drug treatment

#### Cytochalasin D treatment

For fixed cell imaging, parasites invaded confluent HFF monolayer for 4 hours and then treated with 0.2 μM cytochalasin D (CD) or equivalent volume of DMSO for 12 hours before fixation with 4% paraformaldehyde. For live cell imaging, parasites were grown for 12 hours before treatment with 2 μM CD or equivalent volume of DMSO for 30 minutes. Imaging was carried out in the presence of DMSO or CD.

#### IAA Treatment

To deplete TgMyoF protein levels, *TgMyoF-mAID:TgAPT1-EmGFP* parasites were grown in confluent HFF monolayer for 5 or 12 hours and then treated with a final concentration of 500 μM indole acetic acid (IAA) or equivalent volume of 100% ethanol for 6 or 15 hours before fixation for immunohistochemistry or live cell imaging [40,41].

### Creation of transgenic parasite lines

A list of parasite lines, primers and gene accession numbers used in this study can be found in Tables 1, 2 and 3 respectively.

#### Parasite transfection

Parasite transfection was conducted as previously described [42]. Briefly, linearized or circular plasmid DNA or CRISPR HR oligo (produced by PCR) was ethanol precipitated, washed twice 70% ethanol pre-chilled to -20°C and resuspended in cytomix (120mM KCl, 10mM potassium phosphate (pH 7.6), 25mM HEPES (pH 7.6), 2mM EGTA (pH 7.6), 5mM MgCl_2_, 2mM potassium ATP, 150uM CaCl_2_, 5mM reduced glutathione) and electroporated using a BTX electroporator set as follows: voltage 1500 V, capacitance 25 µF and resistance 25 Ω.

#### Creation of TgAPT1-EmGFP parasite line

In the *TgAPT1-EmGFP* parasite line the endogenous TgAPT1 gene is tagged with EmeraldGFP (EmGFP). The pTKOII_APT1EmGFP plasmid was created from pTKOII_MyoF-EmGFP plasmid [39] by digesting with AvrII and BglII restriction enzymes to remove the TgMyoF genomic sequence. 2 kb of genomic DNA upstream of the TgAPT1 stop site was amplified by PCR using Q5 high fidelity polymerase (New England BioLabs) using the Ku80 genomic DNA as a template and the primers TgAPT1 F1 and TgAPT1 R1 (Table 2). The plasmid backbone and the PCR product were gel purified. Ligation of the plasmid was carried out via Gibson assembly using NEB builder HiFi DNA assembly master mix as per manufacturer’s instructions (New England Biolabs; catalog # M5520AA2) and transfected into NEB5α bacteria. Positive clones were screened by colony PCR and verified by sanger sequencing. pTKOII-TgAPT1-EmGFP plasmid was linearized for homologous recombination using the PstI restriction enzyme and 25μg was transfected into 1×10^7^ *ΔKu80: ΔHXGPRT* parental parasites [43,44]. 24 hours after transfection mycophenolic acid in ethanol and Xanthine in 0.3M NaOH were added to growth media to a final concentration of 25μg/ml and 50μg/ml respectively. Drug treatment continued until at least 75% of the parasites were EmGFP positive. Clonal parasites lines were FACS sorted into a single 96-well plate (2 per well) and grown for 7 days to isolate clonal parasite lines resulting in the creation of TgAPT1-EmGFP parasite line. To verify integration of this construct into the APT1 genomic locus, genomic DNA was isolated using Qiagen DNeasy blood and tissue kit as per manufacturer’s instructions (Qiagen, Germantown, MD) and confirmed via PCR using the primers TgAPT1-Promoter F1, APT1 CDS R, EmGFP R (Table 2) (Fig. S1).

#### Creation of TgMyoF-mAID::TgAPT1-EmGFP parasite line

In the *TgMyoF-mAID::TgAPT1-EmGFP* parasite line, the endogenous TgMyoF gene is tagged with HA-mAID [39] and TgAPT1-EmGFP is expressed under the control of the APT1 promoter from the dispensable UPRT locus. For the creation of 5’UPRT-pAPT1-APT1-EmGFP-3’UPRT plasmid (which will be used as a template for created of the HR oligo via PCR), 5’UPRT-pMyoF-mcherry-TgMyoF-3’UPRT (unpublished plasmid) was digested with AvrII and AflII restriction enzyme to remove the pMyoF-mCherry-TgMyoF sequence and replace with pAPT1-TgAPT1-EmGFP fragment, containing 1Kb of upstream of the APT1 start codon and the APT1 coding sequence. TgAPT1-EmGFP sequence was amplified from the pTKOII_APT1-EmGFP plasmid and primers 5’-APT1-UPRT-F and 3’-APT1-UPRT-R (Table 2). The plasmid backbone and the PCR product were gel purified. Ligation of the plasmid was carried out via Gibson assembly as per manufacturer’s instructions (New England Biolabs; catalog # M5520AA2) and transfected into NEB5alpha bacteria. Positive clones were screened by colony PCR and verified by sanger sequencing.

The CRISPR HR oligo was generated by PCR using UPRT F and UPRT R primers and the 5’UPRT-pAPT1-APT1-EmGFP-3’UPRT plasmid as a template with Q5 high fidelity polymerase and 50ul of PCR reaction was cotransfected with 25 μg of pSag1::Cas9::U6::sgUPRT plasmid [45] (AddGene plasmid # 54467) into 1×10^7^ TgMyoF-mAID parasites [39]. 24 hours post-transfection parasites were treated with 10 μM fluorodeoxyribose (FUDR) (which selects for loss of the UPRT gene [46]) until at least 75% of the parasites were EmGFP positive. Parasites were FACS sorted for EmGFP positivity into a single 96-well plate (2 per well) and grown for 7 days. Single clones were selected for further analysis. Genomic DNA was isolated using Qiagen DNeasy blood and tissue kit as per manufacturer’s instructions (Qiagen, Germantown, MD). Genomic DNA was analyzed for correct insertion into the UPRT locus via PCR using primers 5’-UPRT F, APT1 CDS R and UPRT-Intron R (Table 2) (Fig. S2).

#### Creation of TgMyoF-mAID; TgAPT1-Halo parasite line

In the *TgMyoF-mAID:TgAPT1-Halo* parasite line, the endogenous TgMyoF gene is tagged with HA-mAID [39] and TgAPT1-Halo is expressed under the control of the APT1 promoter from the dispensable UPRT locus. For the creation of 5’UPRT-pAPT1-APT1-Halo-3’UPRT plasmid (which will be used as a template for created of the HR oligo via PCR), the 5’UPRT-pAPT1-APT1-EmGFP-3’UPRT plasmid was digested with AflII and BglII to remove the EmGFP coding sequence. The Halo coding sequence was amplified using Halo-F1 and Halo-R2 primers using pmin-ActinCB-Halo as a template (Table 2). The plasmid backbone and the PCR product were gel purified. Ligation of the plasmid was carried out via Gibson assembly as per manufacturer’s instructions (New England Biolabs; catalog # M5520AA2) and transfected into NEB5alpha bacteria. Positive clones were screened by colony PCR and verified by sanger sequencing.

The CRISPR HR oligo was generated by PCR using UPRT F and UPRT R primers and the 5’UPRT-pAPT1-APT1-Halo-3’UPRT plasmid as a template with Q5 high fidelity polymerase and 50ul of PCR reaction was cotransfected with 25 μg of pSag1::Cas9::U6::sgUPRT plasmid [45] (AddGene plasmid # 54467) into 1×10^7^ TgMyoF-mAID parasites [39]. 24 hours post-transfection parasites were treated with 10 μM fluorodeoxyribose (FUDR) (which selects for loss of the UPRT gene [46]) until at least 75% of the parasites were EmGFP positive. Parasites were cloned by limited dilution into a single 96-well plate (2 per well) and grown for 7 days. Single clones were screen for positives by IFA.

#### Creation of ectopic ATG8 expression plasmid

Generation of pAtg8-3xTy-Atg8 plasmid for ectopic expression of TgAtg8 under the endogenous promoter was a 2-step process. First, we created the pAtg8-tdTomato-Atg8 plasmid by digesting the pTub_tdTomato-Atg8 [28] with SfoI and BglII restriction enzymes to remove the tubulin promoter sequence. TgAtg8 promoter was amplified from the Ku80 parental genomic DNA using the primers Atg8 promoter F and Atg8 promoter R. Plasmid backbone and the PCR product was gel purified. Ligation of the plasmid was carried out via Gibson assembly as per manufacturer’s instructions (New England Biolabs; catalog # M5520AA2) and transfected into NEB5alpha bacteria. Positive clones were screened by colony PCR using Atg8 promoter F and Atg8 promoter R (Table 2) and verified by sanger sequencing. In the second step, pAtg8-tdTomato-Atg8 was digested with AvrII and BglII restriction enzymes to remove tdTomato coding sequence. The3xTy tag was created by PCR using primers 3xTy F and 3xTy R (Table 2) and the pTKOII_MyoL-3xTy-mAID plasmid as a template (unpublished plasmid; Table 1). Ligation of the plasmid with 3Xty insert was carried out via Gibson assembly as per manufacturer’s instructions (New England Biolabs; catalog # M5520AA2) and transfected into NEB5alpha bacteria. Positive clones were screened by colony PCR using primers Atg8 promoter F and 3xTy R and verified by sanger sequencing.25 μg of the pAtg8-3xTy-Atg8 plasmid was transiently transfected into the TgMyoF-mAID parasite line and were grown for 12h before treating the parasites with either ethanol or IAA for 6h.

### Fluorescence microscopy

All live and fixed images were acquired on a DeltaVision Elite microscope system built on an Olympus base with a 100x 1.39 NA objective in an environmental chamber heated to 37 ℃. This system is equipped with a 15-bit scientific CMOS camera and DV Insight solid state illumination module with excitation wavelengths DAPI = 390/18nm, FITC = 475/28nm, TRITC = 475/28nm and Alexa 647 = 632/22nm. Single band pass emission filters had the following wavelengths DAPI 435/48nm, FITC = 525/48nm, TRITC = 597/45nm and Alexa 647 = 679/34nm. Image acquisition speeds for the live cell imaging were determined on a case-by-case basis and are indicated in the figure legends. All cells were imaged in multiple focal planes and Z-series were captured at 0.2 μm (fixed) or 0.4 μm (live) steps. Images presented in the figures represent maximum intensity Z-slice projections. Brightness and contrast of images were optimized for print.

### Immunocytochemistry and live cell imaging

Parasites were fixed with freshly made 4% paraformaldehyde (Electron microscopy sciences, Catalog # 15714) in 1x PBS for 15 minutes at room temperature. Cells were washed three times in 1x PBS and permeabilized in 0.25% Triton X-100 (ThermoFisher) in 1x PBS for 15 minutes at room temperature. Cells were washed three times in 1x PBS and were blocked in 2% BSA in 1x PBS and incubated for 15 minutes. All antibodies were diluted in 2% BSA/1x PBS at the concentrations listed in the Table 4. Cells were incubated sequentially with primary and secondary antibody at room temperature for 30 minutes each, with 3 washes with 1x PBS between incubations. To stain DNA parasites were incubated in 10 µM DAPI diluted in 1x PBS for 10 minutes and then washed three times in 1x PBS. Coverslips were mounted onto slides using either Prolong Gold (ThermoFisher Catalog # P36930) or Prolong diamond (ThermoFisher Catalog # P36970) anti-fade reagent and allowed to dry overnight before imaging.

For live cell imaging, growth media was replaced with Fluorobrite DMEM (ThermoFisher) containing 1% FBS and 1x antimycotic/antibiotic pre-warmed to 37 ℃.

The halo pulse chase assay was carried out as per manufacturer’s instructions (Promega, Inc). Parasites were grown overnight in confluent HFF monolayers in MetTak dishes. Parasites were incubated for 15 minutes with 200nM Halo-ligand JF646 diluted in growth media. Cells were wash three times in growth media and incubated for 30 minutes. Cells were washed again two times in growth media and incubated for 5.5 hours. Cells were than labeled in 5µM Halo-ligand TMR for 15 minutes. Cells were wash three times in growth media and incubated for a further 30 minutes in growth media. Cells were washed twice in Fluorobrite DMEM with 1% FBS before imaging cells live.

### Image analysis

#### Quantification of TgAPT1 Vesicle number

To determine the number of APT1 vesicles during parasite division. The apicoplasts were categorized according to their morphology to distinct stages of apicoplast division stage (interphase, elongation, and post fission) by visualizing apicoplast morphology with anti-Cpn60 IFA and expression of TgAPT1-EmGFP. Cell cycle stage by determined by expressing ptub-mCherry-TubulinA1 a marker for the pellicle of daughter and mother parasites. Number of vesicles were counted manually on average intensity projection images using cell counter plugin in Fiji (Fiji is Just ImageJ; http://imageJ.net). Three independent experiments were performed with 25 parasites per experiment counted for each stage of division.

#### Characterization of TgAPT1 vesicle motion

To characterize TgAPT1 vesicle motion, vesicle tracking analysis was performed using Imaris v9.9.0 microscopy image analysis software (Bitplane AG, Oxford Instruments). Z-stack movies that were bleach corrected in ImageJ using the histogram matching algorithm were uploaded to Imaris and brightness and contrast was auto-adjusted using the “normalize time point” extension. Movies were cropped to remove areas that did not contain apicoplast and vesicles. To prevent aberrant tracking inside the apicoplast itself, the apicoplasts were demarcated from vesicles using the surface tool with a lower surface threshold of 1500 arbitrary units (a.u.). Vesicles were identified using the spots creation wizard. Size parameter (estimated XY diameter) was 0.3 µm. To distinguish between vesicles and background fluorescence, only objects with a fluorescence intensity above 150 a.u. were tracked. Tracks were created using autoregressive motion algorithm. To be considered true vesicles, vesicles must be located within 5 µm from the apicoplast surface, to prevent the aberrant detection of fluorescent puncta outside the parasite, especially at later time points when photobleaching reduced the fluorescence intensity of the vesicles. A maximum gap of 2 frames was permitted, (i.e., maximum number of the consecutive time points that a track segment can be missed to be considered continuous). Tracks with less than 5 data points were excluded from the analysis as it was not possible to accurately assign a straightness value. Two independent experiments with a total of 14-17 videos were used for this analysis. Straightness values were calculated by determining the ratio of track displacement (the distance between first and last positions of the vesicle) to track length (total distance traveled).

#### Measuring apicoplast to centrosome distance

Distance between the elongated apicoplast and centrosomes was determined using the straight-line tool in ImageJ by measuring the closest distance from each centrosome to the apicoplast. The distance was considered as 0 when there was an overlap between fluorescence pixels in each channel. Three independent experiments were performed (N=108 and 93 from control and knockdown respectively).

#### Measuring apicoplast length

Elongated apicoplast length was measured using segmented/straight line tool in Fiji. Three independent experiments were performed (N=60 parasites total).

#### Daughter cell-apicoplast proximity categories

For determining the apicoplast proximity to growing daughter cells, we visualized apicoplast localization (Cpn60 staining) in growing daughter parasites (based on eGFP-tubulin). Daughter proximity to the apicoplast was determined visually and categorized as being near to both the daughters (2D), near to one of the daughters (1D) or not close to either daughter (none). Three independent experiments were performed (N=60 total).

#### TgAtg8 localization

TgAtg8 localization during different stages of apicoplast division was determined using anti-Ty antibody staining (kind donation of Dr. Chris de Graffenried: Brown University). Two independent experiments were performed (25 parasites per experiment).

### Statistical analysis

Statistical analysis was performed using GraphPad Prism software. Statistics on data sets were performed using unpaired t-tests. P-values <0.05 were considered statistically significant. Superplots were made as described in Lord et al., [47].

## RESULTS

### The number of TgAPT1-EmGFP vesicles in the cytosol increases during apicoplast elongation

To investigate the mechanisms of apicoplast division and apicoplast protein trafficking from the ER, we generated a parasite line where the endogenous TgAPT1 gene was tagged with EmGFP (Emerald green fluorescent protein) (Fig. 1A and 1B; Fig S1) as TgAPT1 localizes to vesicles in the parasite cytosol and the apicoplast [14,16,48]. Previously published data indicated that APT1 vesicles are formed in a cell cycle dependent manner so we transiently expressed mCherry-TubulinA1 which demarcates the parasite periphery of both the mother and growing daughter parasites to enable quantification of vesicle number at each stage of the parasite cell division cycle. Parasites were also stained with an anti-Cpn60 antibody, a marker of the apicoplast lumen. Based on the apicoplast shape and the presence and size of daughters, parasites were categorized as interphase (no daughter parasites, circular apicoplast), elongating (small daughters, elongated apicoplast) or post-fission (large daughters, circular apicoplast in each daughter) and the number of vesicles per parasite was determined at each cell cycle stage. In interphase parasites, TgAPT1 forms a ring that surrounds the TgCpn60 signal consistent with its localization at the apicoplast membrane (Fig. 1B) and the number of TgAPT1 positive (TgAPT1+) vesicles per parasite is low (1.15±0.07, mean±SEM) (Fig. 1B and 1C). During elongation, the number of TgAPT1 vesicles in the cytosol increased 7-fold to 8.08±0.5 vesicles per parasite (Fig. 1B and 1C). After apicoplast fission and segregation into the daughter cells, the number of TgAPT1 vesicles per parasite decreased to 3.68±0.2 (Fig. 1B and 1C). Notably, Cpn60, a nuclear encoded protein that localizes to the apicoplast lumen, was not found in TgAPT1+ vesicles as previously reported [48] (Fig. 1B, inset). Consistent with the observations and interpretations made by [14], we hypothesize that these vesicles are ER-derived and transport newly synthesized proteins to the apicoplast in a cell cycle dependent manner.

To further investigate the origin of the APT1 vesicles, and determine if these vesicles are derived from or trafficking to the apicoplast, we designed a fluorescent pulse chase experiment using a parasite line expressing APT1-Halo under the endogenous APT1 promoter. Parasites were grown overnight and then pulse labeled with Halo-Ligand JF646 for 15 minutes (Fig. 1D). After this “pulse” labeling parasites were grown without ligand for 6 hours, labeled with Halo-Ligand TMR for 15 minutes (chase) and imaged immediately. Interphase parasites containing a circular apicoplast were labeled with JF646 only indicating that proteins synthesis did not occur after the first pulse labeling. In contrast, during apicoplast division, indicated by elongated apicoplasts, the apicoplast was labeled with both JF646 and TMR. In addition, Halo-Ligand TMR labeled newly synthesized protein that localized to the ER/nuclear envelope and accumulated at the apical and basal ends of the nucleus (Fig. 1D; green arrowheads). We did not observe Halo-JF646 labeled vesicles in the cytosol during apicoplast elongation, as would be expected if the APT1 vesicles originated at the apicoplast. Unfortunately, due to the prevalent ER localization of Halo-TMR it was not possible to discern if TMR labeled vesicles were also present in the cell as would be expected if the APT1 vesicles were derived from the ER. Notably, the ER localization of APT1 was never observed in the APT1-EmGFP construct, likely due to inefficient fluorescent protein maturation in the ER. Collectively, these results show that APT1 protein synthesis and the appearance of the APT1 vesicles occur in a cell cycle dependent manner.

### Actin depolymerization and TgMyoF knockdown leads to an increased number of TgAPT1-EmGFP vesicles in the cytoplasm

TgAct1 and TgMyoF are cytoskeletal proteins with previously established roles in vesicle transport [38,39]. Therefore, we wanted to determine if these proteins play a role in vesicle trafficking to the apicoplast. To do this a previously generated auxin inducible TgMyoF knockdown parasite line TgMyoF-mAID [39] was modified to express TgAPT1-EmGFP under the endogenous TgAPT1 promoter from the dispensable UPRT locus to create TgMyoF-mAID::TgAPT1-GFP parasite line (Fig. S2A).

To knockdown TgMyoF, TgMyoF-mAID::TgAPT1-EmGFP parasites were treated for 15 hours with ethanol (control) or 500 μM IAA, which results in depletion of TgMyoF to undetectable levels after 4 hours [39]. To depolymerize actin, TgAPT1-EmGFP parasites were treated with DMSO (control) or 0.2 µM cytochalasin D (CD) for 12h prior to fixation. Parasites transiently expressing mCherry-Tubulin were categorized as interphase, elongation or post-fission as described for figure 1. Both actin depolymerization and TgMyoF depletion resulted in an approximately 2-fold increase in the number of TgAPT1-EmGFP positive vesicles at all stages of the apicoplast division cycle (Fig. 2A-F; Fig. S3 & S4). In both treatment conditions, an apicoplast inheritance defect was observed as expected [37] (Fig. 2A&2D; magenta arrow). “Apicoplast-less” parasites contained the highest accumulation of vesicles in comparison to all other stages of apicoplast division (Fig. 2C and F).

**Figure 2.**
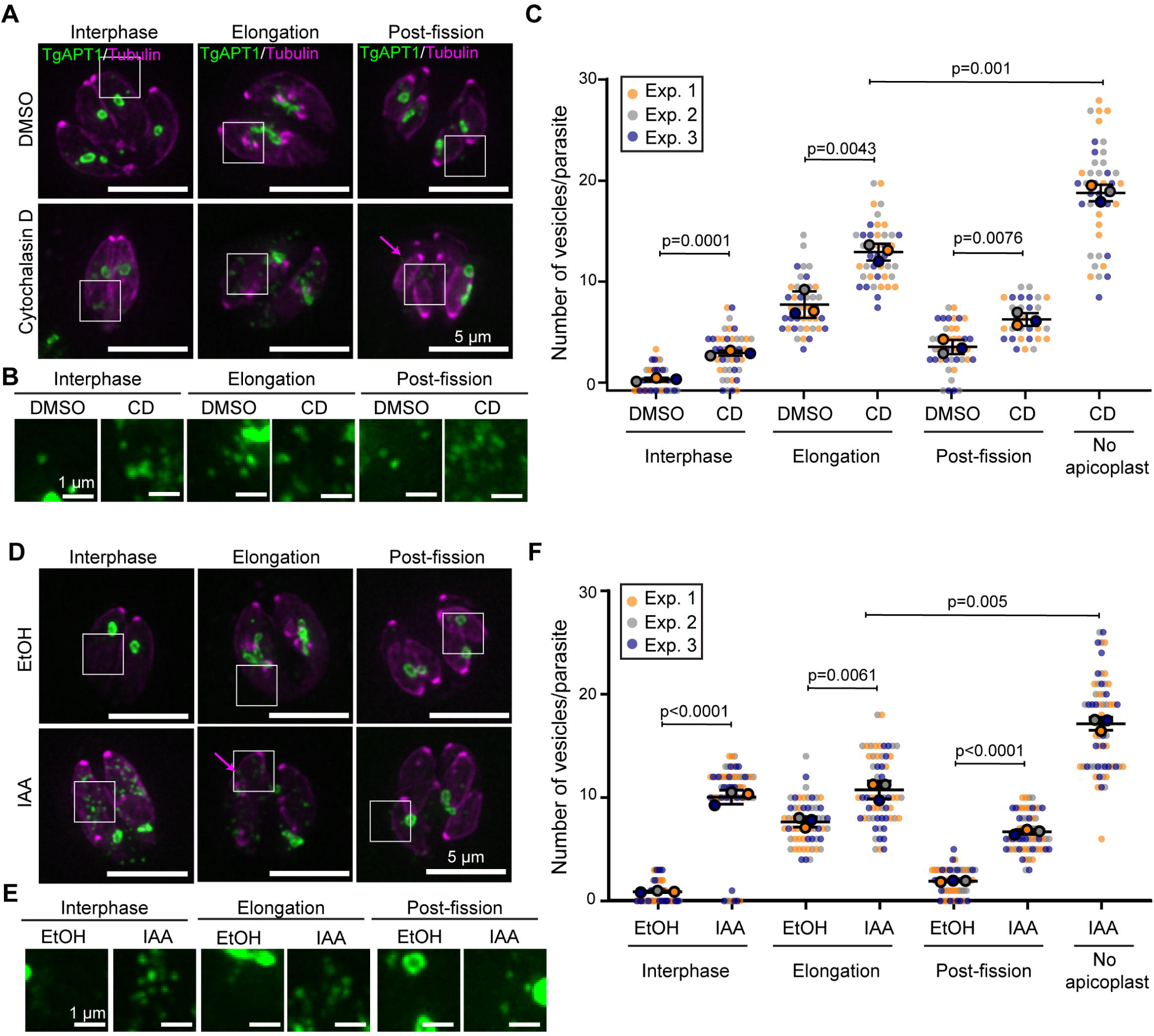
Actin depolymerization and TgMyoF knockdown results in an accumulation of APT1 vesicles. (A) TgAPT1-EmGFP parasites treated with DMSO (control) or 0.2 μM cytochalasin D for 12 hours were categorized as either interphase, elongation or post-fission based on the stage of the apicoplast division cycle and the size and presence of daughter parasites. Magenta arrow indicates parasite that does not contain an apicoplast. White box indicates area used to make inset in panel B. Images are maximum intensity projections of deconvolved images. Scale bar represents 5 μm. (B) Inset panels from (A). Brightness and contrast of insets was adjusted to facilitate visualization of dimmer TgAPT1 vesicles. Inset scale bar represents 1 μm. (C) Quantification of the number of TgAPT1-EmGFP vesicles per parasite during interphase, elongation and post-fission stages of the apicoplast division cycle. Parasites with no apicoplast were categorized separately. DMSO: n= 42/apicoplast division stage, n= 126 total; CD: n= 30-42/apicoplast division stage, n = 156 total. TgAPT1 vesicle number per parasite are as follows (Mean±SEM): DMSO - interphase: 1.05±0.1; elongation: 8.26±0.73; and post-fission: 4.19±0.39. CD interphase: 3.59±0.15; elongation: 13.30±0.46; post-fission: 6.83±0.35; no apicoplast: 19±0.46). The mean from each experiment is indicated with large circles. These values were used to calculate the mean, standard error of the mean and p-value. Raw data is shown with smaller colored circles. Experiment 1 in orange, experiment 2 in grey, experiment 3 in blue. Statistical significance was determined by unpaired Student’s *t* test. (D) TgMyoF-mAID:TgAPT1-EmGFP parasites treated with ethanol (control) or 500 μM IAA for 15 hours were categorized as either interphase, elongation or post-fission based on the stage of the apicoplast division cycle and the size and presence of daughter parasites. Magenta arrow indicates parasite that does not contain an apicoplast. White box indicates area used to make inset in panel E. Images are maximum intensity projections of deconvolved images. Scale bar represents 5 μm. (E) Inset panels from (D). Brightness and contrast of insets was adjusted to facilitate visualization of dimmer TgAPT1 vesicles. Inset scale bar represents 1 μm. (F) Quantification of the number of TgAPT1-EmGFP vesicles per parasite during interphase, elongation and post-fission in control and TgMyoF knockdown parasites. Parasites with no apicoplast were categorized separately. (EtOH: n= 60/apicoplast division stage, n= 180 total; IAA: n= 60/apicoplast division stage, n = 240 total). TgAPT1 vesicle number per parasite are as follows (Mean±SEM): Ethanol - interphase: 0.8±0.14; elongation: 7.57±0.27; and post-fission: 1.83±0.16. IAA interphase: 9.9±0.52; elongation: 10.7±0.43; post-fission: 6.6±0.24; no apicoplast = 17.1±0.58). The mean from each experiment is indicated with large circles. These values were used to calculate the mean, standard error of the mean and p-value. Raw data is shown with smaller colored circles. Experiment 1 in orange, experiment 2 in grey, experiment 3 in blue. Statistical significance was determined by unpaired Student’s *t* test.

### TgAPT1 vesicles exhibit predominately diffusive-like motion

Next, we sought to describe the motion characteristics of TgAPT1+ vesicles, as previous work demonstrated that other vesicle types are transported throughout the parasite cytosol in an actin and TgMyoF dependent manner [38,39]. TgAPT1-EmGFP parasites in the elongation phase of apicoplast division were imaged by live cell microscopy. Visual inspection of these recordings revealed two striking observations. First, TgAPT1-EmGFP vesicles were found in the parasite cytosol, confirming observations from the fixed cell experiments (Fig. 1 and 2). The motion of these vesicles was predominately diffusive (Fig 3A; inset 2), although a small number of vesicles exhibited directed motion, i.e., – sustained movement in a single direction for continuous frames (Fig. 3A; inset 1) (Video S1). Secondly, elongated apicoplasts were not static, linear structures, but exhibited changes in morphology throughout the 2-5min imaging period (Fig. 5; Video S2).

**Figure 3.**
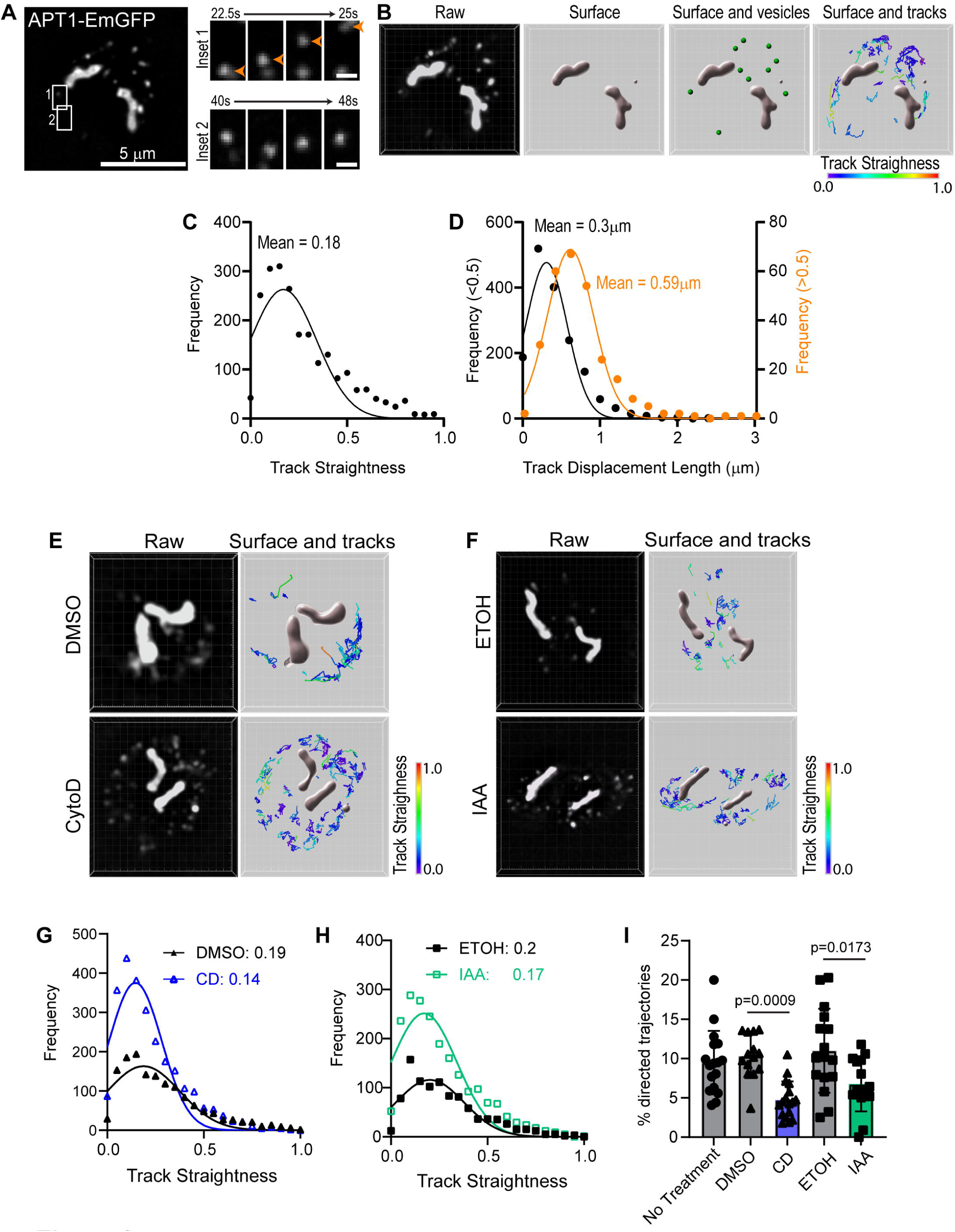
APT1 vesicles exhibit predominately diffusive motion. (A) Localization TgAPT1-EmGFP parasites in live parasites. TgAPT1 is localized to both the apicoplast and vesicles. White boxes indicate areas used to make insets 1 and 2. Inset 1 depicts TgAPT1+ vesicle moving in a directed manner. Inset 2 depicts a TgAPT1+ vesicle moving with diffusive-like movement. Scale bar = 5µM. Inset scale bar = 1 µM. (B) The Z-stack movie of TgAPT1-EmGFP expressing parasites were imported into Imaris. Raw data from frame 1 of the movie is shown. Isosurfaces of the apicoplast were reconstructed to prevent aberrant tracking inside the apicoplast itself. Vesicle tracks observed in the first 20 seconds of imaging are color-coded by straightness value. (C) Frequency distribution of trajectory straightness values of TgAPT1 vesicles. Straightness value close to zero indicates random motion. Straightness value close to 1 indicates directed/linear movement. Mean straightness value is 0.18±0.15. (D) Frequency distribution of displacement lengths of TgAPT1+ vesicles. Vesicles with a straightness value less than 0.5 (black) had a mean displacement length of 0.3±0.27µm (Mean±SD), while vesicles with a straightness value greater than 0.5 (orange) had a mean displacement length of 0.59±0.3µm (Mean±SD). (E) Localization TgAPT1-EmGFP parasites in live parasites treated with DMSO or CD for 30 minutes prior to imaging. Raw data from frame 1 of the movie is shown. Z-stack movies were imported into Imaris, isosurfaces of the apicoplast and vesicles were reconstructed. Vesicle tracks observed in the first 20 seconds of imaging are color-coded by straightness value. (F) Localization TgAPT1-EmGFP parasites in live parasites treated with ethanol or IAA for 15 hours prior to imaging. Raw data from frame 1 of the movie is shown. Z-stack movies were imported into Imaris, isosurfaces of the apicoplast and vesicles were reconstructed. Vesicle tracks observed in the first 20 seconds of imaging are color-coded by straightness value. (G) Frequency distribution of trajectory straightness values of TgAPT1 vesicles from DMSO treated (black) and cytochalasin D (CD) treated (blue) parasites. Mean straightness values are for DMSO and CD treated are 0.19±0.18 and 0.14±0.13 (Mean±SD) respectively. Straightness value close to zero indicates random motion. Straightness value close to 1 indicates directed movement. (H) Frequency distribution of trajectory straightness values of APT1 vesicles from ethanol (black) and IAA (green) treated parasites. Mean straightness values are for ethanol and IAA treated are 0.2±0.18 and 0.17±0.17 (Mean±SD) respectively. Straightness value close to zero indicates random motion. Straightness value close to 1 indicates directed movement. (I) Bar graph indicating the percentage of vesicles exhibiting directed motion, defined as a straightness value greater than 0.5 and a track displacement length of greater than 500nm. Actin depolymerization with CD or TgMyoF knockdown with IAA results in a significant (∼50%) decrease in the percentage of directed trajectories.

**Figure 4.**
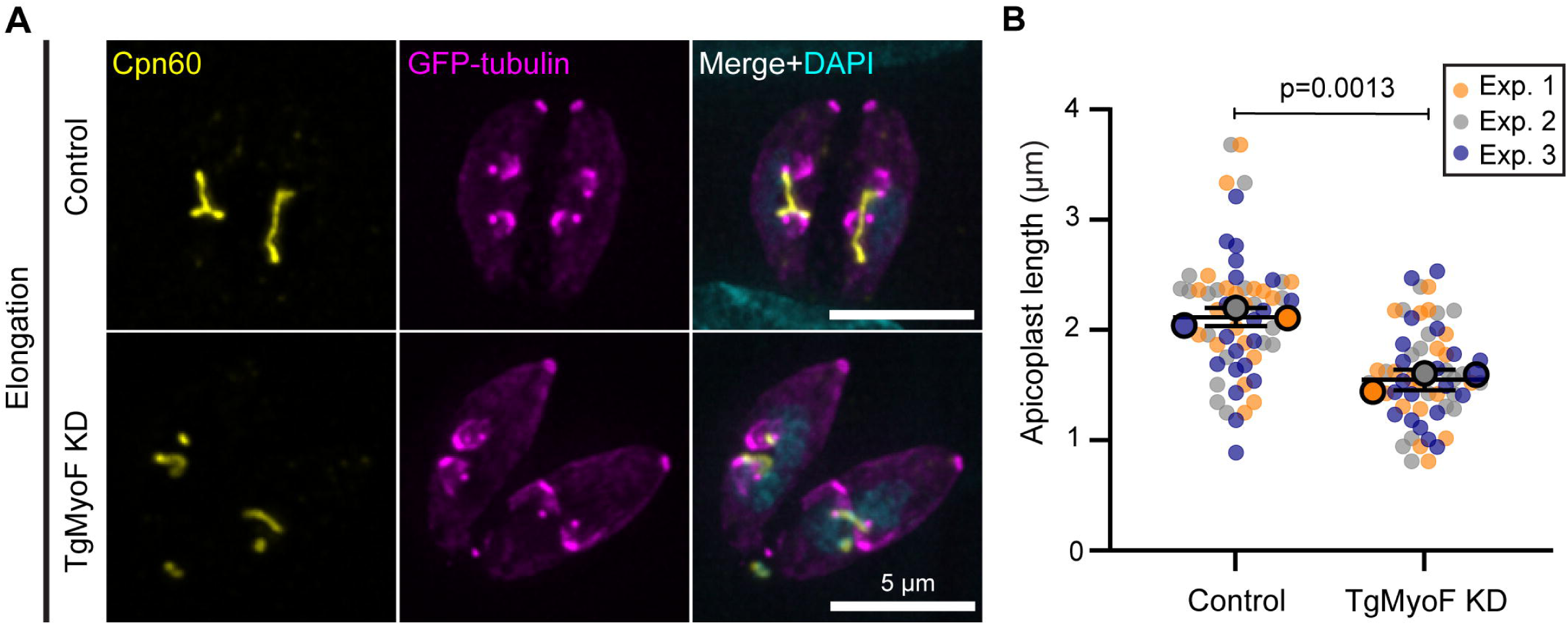
TgMyoF knockdown results in reduced apicoplast length. (A) TgMyoF-mAID parasites expressing eGFP-tubulin (magenta) were grown for 12 hours before treatment with ethanol (control) or IAA (TgMyoF KD) for 6 hours. Parasites were stained with an anti-Cpn60 antibody (yellow) and DAPI (cyan). Scale bar represents 5 μm. (B) Quantification of the apicoplast length in control and TgMyoF KD parasites from 3 independent experiments (20 parasites per experiment). Mean apicoplast length was 2.12±0.05 and 1.55±0.05 µm for ethanol and IAA treated parasites respectively. The mean from each experiment is indicated with large circles. These values were used to calculate the mean, SEM and p-value. Raw data is shown with smaller colored circles. Experiment 1 in orange, experiment 2 in grey, experiment 3 in blue. Statistical significance was determined by unpaired Student’s *t* test.

**Figure 5.**
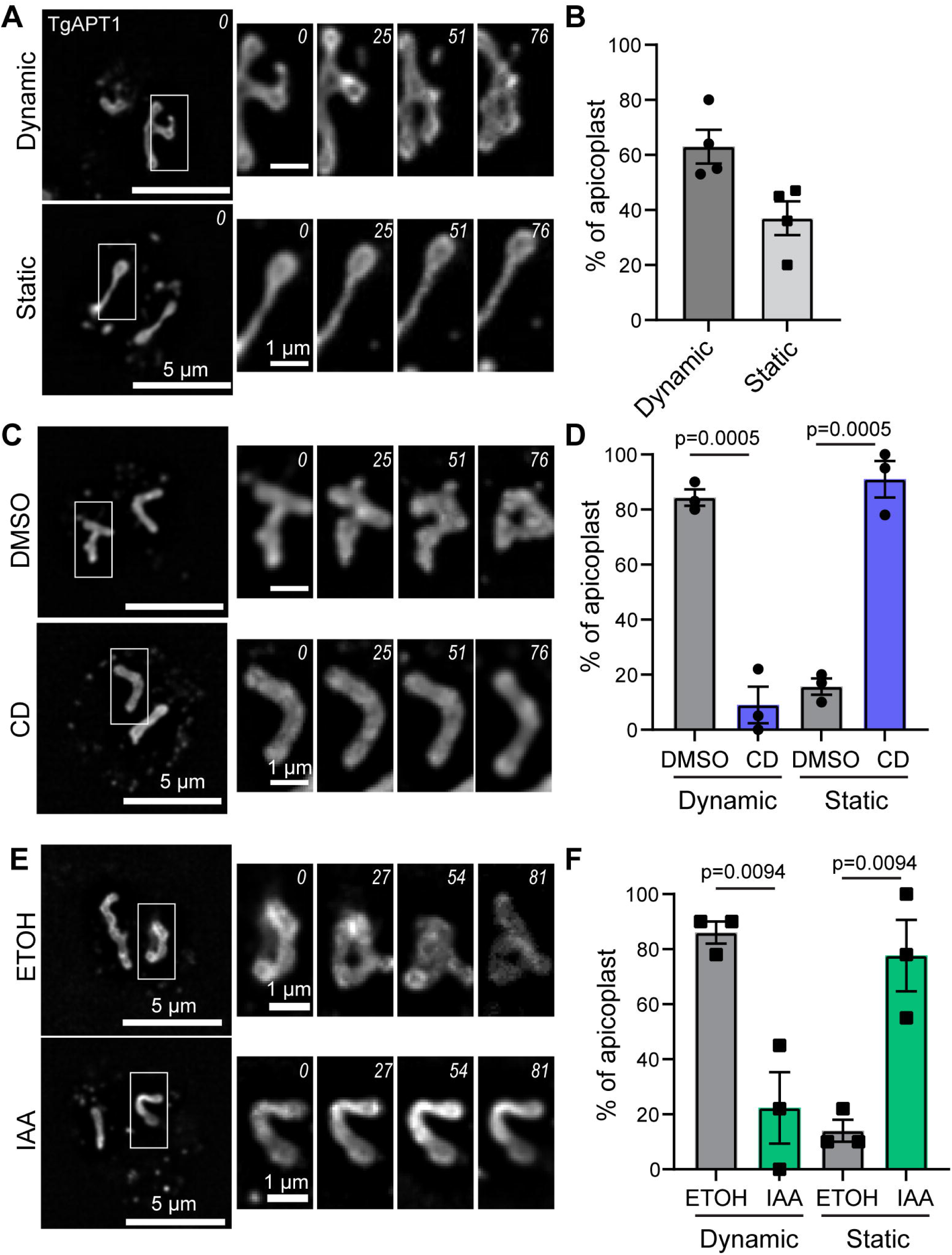
Apicoplast dynamics require actin and TgMyoF. (A) Time lapse images of TgAPT1-EmGFP parasites during the elongation phase of apicoplast division. *Upper:* The apicoplast is dynamic and exhibits branch formation and movement over the ∼2-minute imaging period. *Lower:* An example of a static elongated apicoplast. Apicoplast does not change shape over the imaging period. White box indicates area used to make time-lapse inset. Inset indicates change in apicoplast morphology over time. Time in seconds is indicated in the top right corner of each image. Scale bar represents 5 µm. Inset scale bar = 1 µm. Lower: (B) Quantification of apicoplast dynamics. Apicoplast was categorized as dynamic if one of the following was observed: dramatic changes in apicoplast shape such as bending, branch formation, branch movement or retraction. Apicoplast was categorized as static of there were subtle or no changes in apicoplast shape or position. N=70 from four independent experiments. 63±6.2% of parasites contained dynamic apicoplasts. 37±6.2 contained static apicoplasts. Data is represented as Mean ± SEM. Statistical significance was determined by unpaired Student’s *t* test. (C) Time lapse images of TgAPT1-EmGFP parasites during the elongation phase of apicoplast division imaged after treatment with DMSO or cytochalasin D. White box indicates area used to make time-lapse inset. Inset indicates change in apicoplast morphology over time. Time in seconds is indicated in the top right corner of each image. Scale bar represents 5 µm. Inset scale bar = 1 µm. (D) Quantification of apicoplast dynamics after DMSO or CD treatment. 84±2.9% of DMSO treated parasites (grey) contained dynamic apicoplasts. 9±6.7% of CD treated parasites (blue) contained dynamic apicoplasts. Data is represented as Mean ± SEM. N= 32 and 41 for DMSO and CD respectively from three independent experiments. Statistical significance was determined by unpaired Student’s *t* test. (E) Time lapse images of TgMyoF-mAID:TgAPT1-EmGFP parasites during the elongation phase of apicoplast division imaged after treatment with ethanol or IAA. White box indicates area used to make time-lapse inset. Inset indicates change in apicoplast morphology over time. Time in seconds is indicated in the top right corner of each image. Scale bar represents 5 µm. Inset scale bar = 1 µm. (F) Quantification of apicoplast dynamics after ethanol or IAA treatment. 86±4% of ethanol treated parasites (grey) contained dynamic apicoplasts. 22±13% of IAA treated parasites (green) contained dynamic apicoplasts. Data is represented as Mean ± SEM. N = 38 and 40 for ethanol and IAA respectively. Statistical significance was determined by unpaired Student’s *t* test.

To quantify vesicle movement in an automated and unbiased manner, vesicle motion analysis was performed using the spot creation wizard in Imaris image analysis software (https://imaris.oxinst.com/) (Fig. 3B). To prevent aberrant tracking of fluorescent signal inside the apicoplast itself, a surface mask was created over the apicoplast (Fig 3B; raw vs. surface). Vesicles were automatically detected based on size and fluorescent intensity (Fig. 3B; surface and vesicles) (see materials and methods for details). Vesicles that are transported in an active, motor driven manner typically exhibit “directed trajectories”, i.e.-vesicles move in approximately a straight trajectory [38], while diffusive vesicles exhibit random movements. Thus, to determine the number of vesicles exhibiting diffusive or directed movement, track straightness was calculated by determining the ratio of track displacement (the distance between first and last positions of the vesicle) to track length (total distance traveled). A straightness value close to zero indicates random motion, i.e. – track length is greater than the distance between the first and last positions indicating random motion. A straightness value close to 1 indicates that the total distance travelled and the distance between the first and last positions were similar indicating that the vesicle moved in approximately a straight line. Vesicle tracks observed in the first 20 seconds of imaging are color-coded according to straightness (Fig. 3B). A frequency distribution of the straightness values indicates that over 80% of vesicles have a straightness value less than 0.5 and the mean straightness value was 0.17±0.15 confirming that apicoplast vesicle motion is predominantly diffusive in nature (Fig. 3C). We then compared the track displacement length for trajectories that had a straightness value greater or less than 0.5. Randomly moving vesicles (with a straightness value <0.5) had a mean track displacement value of 0.3µm. While vesicles with straighter trajectories moved twice as far with a mean displacement of 0.59µm (Fig. 3D).

### Loss of TgMyoF and F-actin has only a minor impact TgAPT1-EmGFP vesicle motion

Next, we sought to determine if actin depolymerization or TgMyoF knockdown altered vesicle motion. TgAPT1-EmGFP parasites were grown for 18h prior to treatment with 2 µM CD or equivalent volume of DMSO for 30 min before live cell imaging (Fig. 3E) (Video S3), while to deplete TgMyoF, TgMyoF-mAID::TgAPT1-EmGFP parasites were grown for 15 hours with ethanol or 500µm IAA before live cell imaging (Fig. 3F) (Video S4).

Vesicle movements were again tracked with Imaris as described above. Actin depolymerization or TgMyoF knockdown had no effect on the mean straightness values for DMSO, CD, Ethanol and IAA treated parasites were 0.18±0.18, 0.14±0.13, 0.2±0.17 and 0.17±0.17 respectively (Fig. 3G and 3H). Although, upon loss of F-actin and TgMyoF, the number of vesicles in the cytosol was increased compared to controls. In all cases (treatment and control) the percentage of vesicles with a straightness value <0.5 was ∼80%. To determine if the movement of the remaining ∼20% of vesicles was TgMyoF and actin driven, vesicles with a straightness value greater than 0.5 were analyzed further. Since vesicles with a straightness value with greater than 0.5 exhibited longer track displacement (Mean length was 0.59µm (Fig. 3D)) we defined “directed trajectories” as those with a straightness value >0.5 and a track displacement length of at least 0.5µm. In no treatment, DMSO- and ethanol-treated parasites, the percentage of vesicles which met these criteria were 9.5%, 10.3% and 11% respectively. Upon actin depolymerization with CD or TgMyoF knockdown this percentage was reduced to 4.7% and 6.8% respectively (Fig. 3I). Although these changes are statistically significant, overall our data indicates that only a small portion of APT1 vesicles are transported in active, actomyosin dependent manner. There are two possible origins for these vesicles: those which have bud from the ER and are destined for the apicoplast or vesicles that have originated from the apicoplast.

### TgMyoF knockdown results in reduced apicoplast length

If vesicle accumulation was due to either a defect in protein trafficking to the apicoplast or fragmentation of the apicoplast, we predict that this would alter apicoplast morphology. Therefore, we assessed apicoplast length in the elongation phase of apicoplast division in the absence of TgMyoF. In order to compare parasites at the same stage of cell division, parasites were transiently transfected with a GFP-tubulin expression cassette in order to fluorescently label the pellicle of mother and daughter parasites. Parasites were grown for 12 hours prior to initiating a 6-hour treatment with ethanol or 500 μM IAA. This shortened IAA treatment time compared with previous experiments allowed us to visualize the apicoplast in the first cell division cycle after depletion of TgMyoF. Parasites were then stained with an anti-Cpn60 antibody and DAPI before image acquisition (Fig. 4A). During elongation, TgMyoF knockdown resulted in a ∼25% reduction in apicoplast length (1.54±0.05µm, mean±SEM) when compared to the control parasites (2.11±0.05 µm, mean±SEM) (Fig. 4B). Apicoplast morphology was not disrupted in TgMyoF knockdown parasites in interphase and post-fission stages of division (Fig. S5).

### The apicoplast undergoes morphological changes during elongation

Live cell imaging of TgAPT1-EmGFP parasites demonstrated that the apicoplast morphology in the elongation phase of apicoplast division was highly variable between individual vacuoles. In some vacuoles, the apicoplast appeared to be highly dynamic exhibiting alternating linear, U-shaped and branched morphologies throughout the ∼2-minute imaging period (Fig. 4A; top panel) (Video S2), while others were static in the elongation phase of apicoplast division (Figure 5A; bottom panel) (Video S2). Given this variability, we quantified the percentage of apicoplasts that were highly dynamic and exhibited rapid changes in morphology during the course of imaging (102 seconds) (Fig. 5A and B). ““Dynamic” apicoplasts were defined as those that exhibited one or more of the following morphological changes: extending and/or retracting, bending, or branch formation, or branch movement (Fig. 5A; top panel). 63±6.1% (Mean±SEM) of the apicoplasts were highly dynamic (Fig. 5B). Whereas 37±6.1% (Mean±SEM) of parasites contained either less dynamic apicoplasts, exhibiting subtle changes in morphology, or were static with no observable changes in the morphology throughout the ∼2-minute imaging time (Fig 5A and B).

### Apicoplast dynamics requires actin and TgMyoF proteins

Considering the dynamic nature of the apicoplast, we wanted to determine if these dynamics are dependent on actin and TgMyoF. To depolymerize actin, TgAPT1-EmGFP parasites were again treated with 2 µM cytochalasin D or equivalent volume of DMSO for 30 min before live cell imaging (Fig. 5C). 82±2.9% (Mean±SEM) of the apicoplasts in control DMSO-treated parasites were highly dynamic, while only 9±6.6% (Mean±SEM) of the apicoplasts in the cytochalasin D treated parasites were highly dynamic (Fig. 5D). To test the role of TgMyoF in apicoplast dynamics, TgMyoF-mAID; TgAPT1-EmGFP parasites treated with 500 μM indole acetic acid (IAA) to induce TgMyoF knockdown for 15h prior to live cell imaging (Fig. 5E). Similar to the cytochalasin D treatment, TgMyoF knockdown resulted in reduced apicoplast dynamics. The percentage of parasites with a highly dynamic apicoplast was significantly reduced from 86±4% (Mean±SEM) in control parasites to 22±13% in the TgMyoF KD parasites (Fig. 5F).

### The apicoplast dynamically associates with static centrosomes

Apicoplast association with the centrosomes is a crucial step in the apicoplast division cycle. In the first step in the cell division cycle, the centrosome migrates to the basal end of the nucleus and duplicates, then both centrosomes return to the apical side of the nucleus where they associate with the apicoplast [27,49]. The dynamics of the elongated apicoplast (Fig. 5) have not previously been described and it is unclear if or how the apicoplast interaction with the centrosomes, which are embedded in the nuclear envelope [50], is maintained during apicoplast movement.

To further explore this phenomenon, wild type parasites were transiently transfected with plasmids containing FNR-RFP (a marker for the apicoplast; [27]) and Centrin 1-GFP (a marker for the centrosome [51]) expression cassettes and visualized using live-cell microscopy. In parasites with highly dynamic, elongated apicoplasts the centrosomes were not always associated with the tips of the apicoplast but were frequently associated with the sides of the elongated apicoplast (Fig. 6A; inset 1 and 3) (Video S5). The apicoplast appears to slide in relation to the static centrosome (Fig. 6A, Inset 1, Video S6). The apicoplast was observed to undergo branching when the centrosome was associated with the side rather than the tip of the apicoplast (Fig. 6A; Inset 3, magenta arrow) (Video S7). The most dramatic morphological changes occurred when the apicoplast was associated with only a single centrosome (Fig. 6A, Inset 2). The apicoplast was observed extending towards the second centrosome before retracting backwards. In contrast to the dynamics observed in the elongation phase of apicoplast division, interphase and post-fission apicoplasts were static. In interphase parasites the apicoplast was circular and did not change shape over the course of imaging (Fig. 6B; upper panel) (Video S8). After apicoplast fission, the apicoplast remains elongated for a time before retraction into a circular morphology. Post-fission elongated apicoplasts are static over the course of the imaging period. The apicoplast movement and branching that were observed prior to fission are no longer observed and centrosomes are statically associated with the apicoplast tips (Fig. 6B; lower panel) (Videos S9).

**Figure 6:**
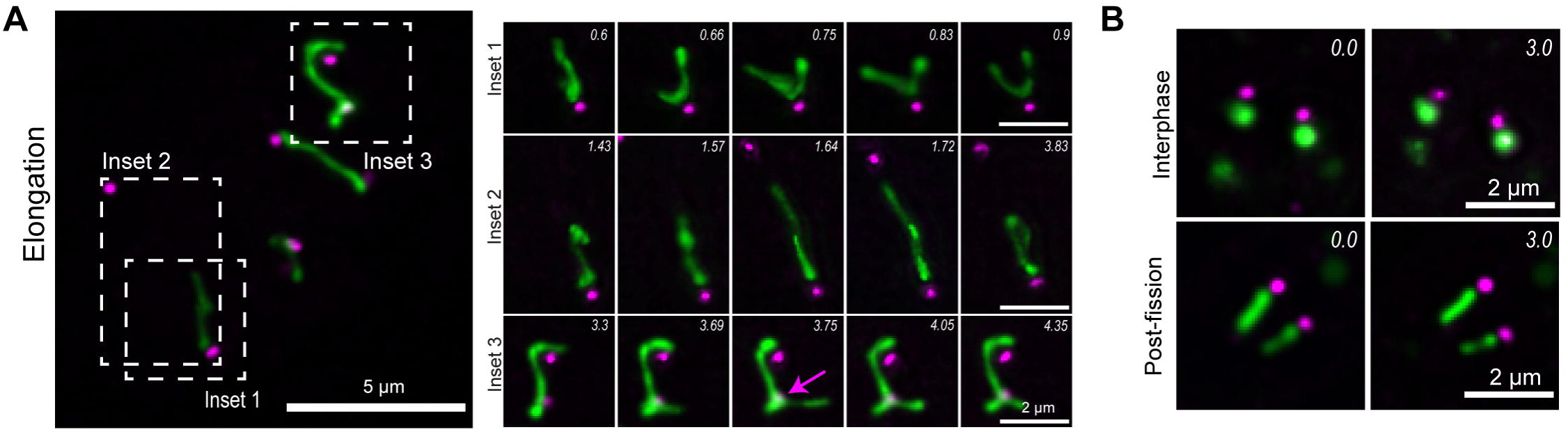
Centrosome-apicoplast interaction is maintained during apicoplast dynamics. (A) Maximum intensity projection of a deconvolved images of parasites expressing eGFP-centrin1 (centrosome marker; magenta) and FNR-RFP (apicoplast marker; green) during the elongation phase of apicoplast division. Parasites were imaged on 5 focal planes for a total of 10 minutes. Z-slice step size was 0.4 µm. Dashed boxes indicate areas used to make time lapse insets. Inset 1: The apicoplast is attached to a single centrosome and slides in relation to the static centrosome, associating with both the tips and sides of the apicoplast. Inset 2: The same apicoplast at a later time point undergoes dramatic morphological change, extending to ∼3 μm in length towards the second centrosome before collapsing backward. Inset 3: Two centrosomes associated with the sides of the apicoplast. Apicoplast “branching” (magenta arrow) is observed at the site of centrosome association. Imaging time points in minutes are indicated in the top right-hand corner of each image. Scale bar represents 5 µm, inset scale bar represents 2 μm. (B) Interphase (upper) and post-fission (lower) apicoplasts did not exhibit morphological changes throughout the course of imaging. Imaging time points in minutes are indicated in the top right-hand corner of each image. Scale bar represents 2 µm.

### TgMyoF knockdown disrupts apicoplast and centrosome interaction

The live cell imaging of apicoplast and centrosome interactions, suggest that the actomyosin-driven apicoplast dynamics are important for initiating the interaction between these two structures and positioning of the centrosomes at the apicoplast tips (Fig. 6). Therefore, we directly investigated if the loss of TgMyoF, which disrupted apicoplast dynamics (Fig. 5), also disrupted apicoplast-centrosome interactions. TgMyoF-mAID parasites transiently expressing the centrosome marker Centrin1-GFP were grown for 12 hours post invasion and then treated with ethanol or 500 μM IAA for 6 hours before fixation and staining with an anti-Cpn60 antibody and DAPI. In control parasites, at the early stages of apicoplast division when the length of the elongated apicoplast is short (less than 1.5µm), the centrosomes and the apicoplast are frequently not associated with one another (Fig. 7A) and the mean apicoplast to centrosome distance is 0.99±0.18µm (Mean±SEM) (Fig. 7B&C). Later in the apicoplast division cycle, when the length of the apicoplast is greater than 1.5µm, the mean centrosome-apicoplast distance is significantly shorter at only 0.36±0.05µm (Fig. 7A-C) and the centrosomes were close with either the sides or the tips of the apicoplast (Fig. 7A). However, in TgMyoF knockdown parasites the apicoplast to centrosome distances were 1.2±0.12 and 1.0±0.09 µm when the apicoplast length was greater than or less than 1.5µm respectively. (Fig. 7A-C) indicating that TgMyoF is needed for recruiting the apicoplast to the centrosomes.

**Figure 7.**
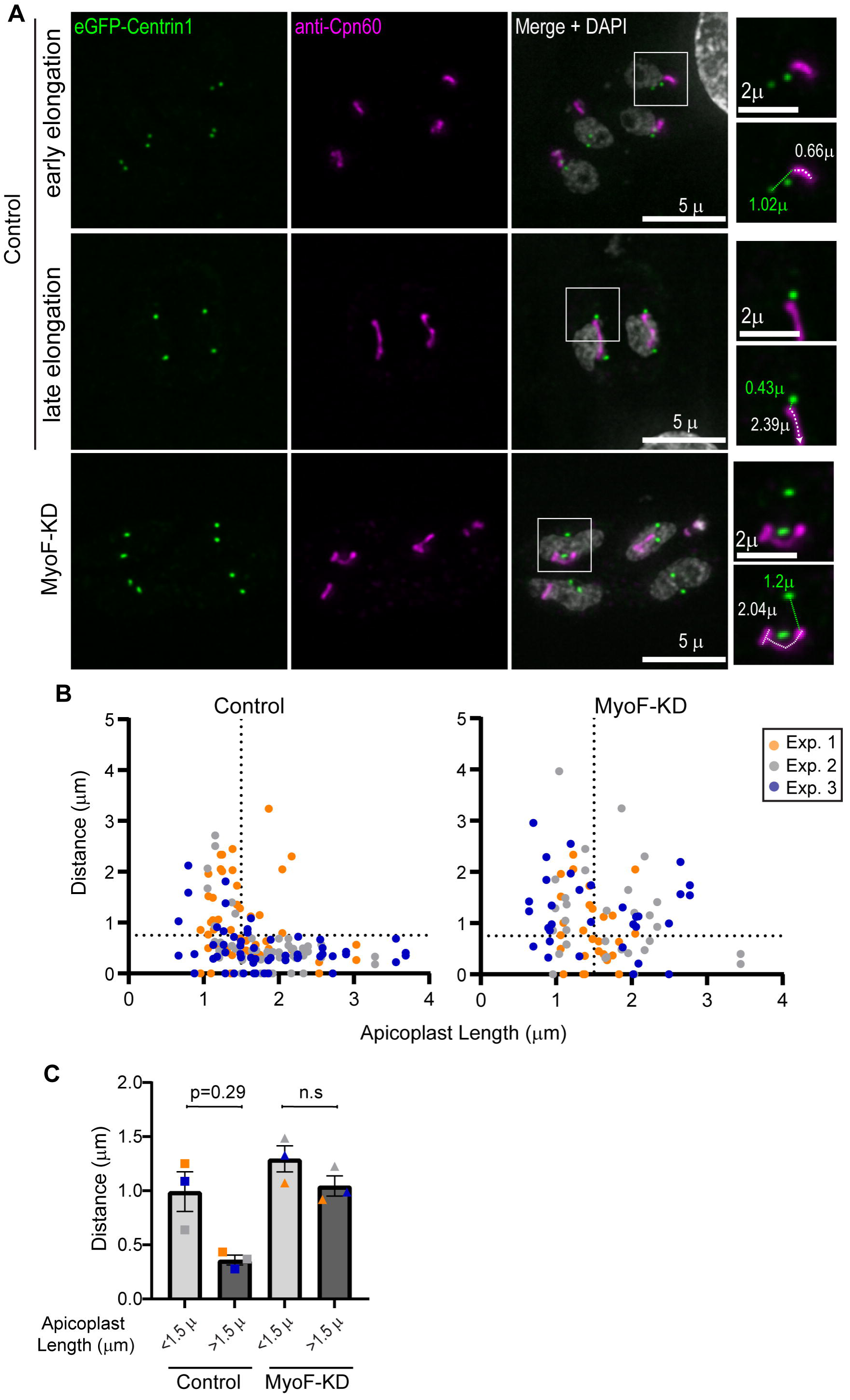
TgMyoF knockdown disrupts the apicoplast and centrosome interaction. (A) TgMyoF-mAID parasites expressing eGFP-Centrin1 (Green) were grown for 18h before treatment with either ethanol or IAA for 6h. Parasites were fixed and stained with an anti-Cpn60 (magenta) antibody and DAPI (grey). Images are maximum intensity projections of deconvolved images. Based on the length of the apicoplast control parasites were categorized as in either early or late elongation. Since the apicoplast elongation length was reduced (Fig. 4) in TgMyoF-KD parasites, these parasites could not be categorized into early or late stages of elongation. White box in indicates area used to make inset. Inset: White dashed line and white number indicate apicoplast length. Green dashed line and green number indicate the distance between the apicoplast and the centrosome. Scale bar represents 5 µm. Inset scale bar represents 2 µm. (B) Distance between the centrosome and apicoplast were plotted as a function of apicoplast length in control and MyoF-KD parasites. Experiment 1 in orange, experiment 2 in grey, experiment 3 in blue. In control parasites, the apicoplast-centrosome distance is short when apicoplast length is above 1.5µm. (C) Parasites were categorized has having an apicoplast greater or less than 1.5µm and the mean apicoplast-centrosome distance was calculated for each category in control and TgMyoF-KD parasites. In control parasites, the apicoplast centrosome distance is significantly shorter in parasites containing a long apicoplast (>1.5µm) compared with control parasites containing a short apicoplast (<1.5µm). In TgMyoF-KD parasites the mean apicoplast-centrosome distance was ∼1µm, regardless of apicoplast length.

We next assessed how the loss of centrosome interaction affected apicoplast inheritance. In 100% of control parasites, elongated apicoplasts were found to be in close proximity to both of the growing daughter parasites. In early stages of division, the apicoplast was adjacent to the daughter parasites (Fig. 8A; white arrow head). Later in division, both daughters contain one end of the elongated apicoplast, even when one the daughters was in the “upside down” orientation, (i.e. the apical ends of the daughter parasites have opposite orientations within the mother) (Fig 7C; magenta arrowhead). Upon depletion of TgMyoF only 13.3% of elongated apicoplasts were in close proximity to both the growing daughter parasites (2D) while 70% of the elongated apicoplasts were in close proximity to only one of the daughter parasites (1D) (Fig. 8A; white arrow), and 16.6% of the elongated apicoplasts were not found to be in close proximity to either of the growing daughter parasites (none) (Fig. 8B and Fig. S6). Thus, the disruption of the centrosome-apicoplast interaction leads to asymmetric segregation of the apicoplast into a single daughter cell or to the residual body.

**Figure 8:**
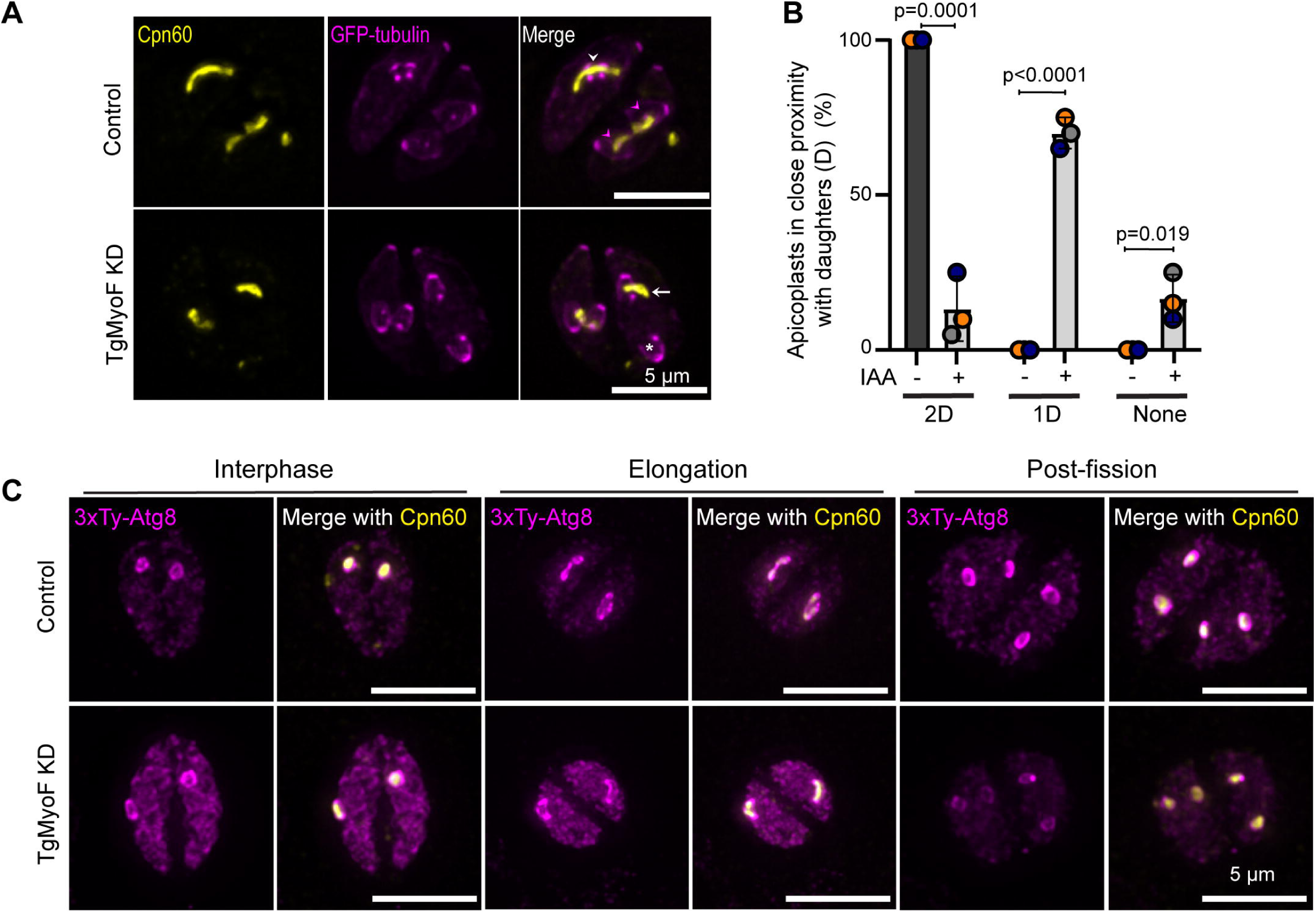
Apicoplast inheritance defects are not due to changes in Atg8 localization. (A) TgMyoF-AID parasites expressing eGFP-tubulin (magenta) were treated with ethanol or IAA for 6 hours then fixed and stained with an anti-Cpn60 antibody (yellow). Control parasites show that the elongated apicoplasts are in proximity to the respective daughter parasites (white arrowheads). Later in division as indicated by larger daughter parasites, the tips of the elongated apicoplast are localized within the growing daughters (magenta arrowheads). Upon TgMyoF knockdown, there is an asymmetric localization of the elongated apicoplast to one of the daughter cells (white arrow) and absent from second daughter parasite (asterisks). Scale bar represents 5 μm. (B) Quantification of the percentage of daughter parasites in close proximity to the elongated apicoplast in control parasites and TgMyoF KD parasites. Parasites were categorized as follows: 2D – elongated apicoplast near or within both daughter parasites. 1D -elongated apicoplast in close proximity to one of the daughter parasites, and “none” - elongated apicoplast not in close proximity to either of the daughter parasites. In control parasites, 100% of apicoplast were in close proximity to both daughter parasites compared to 13.33±6% of TgMyoF-KD parasites. In TgMyoF-KD parasites, 70±2.9% and 16.7±4.4% of apicoplasts were associated with one daughter parasite or neither daughter parasite respectively (Mean±SEM). Bar graph shows quantification of combined data from 3 independent experiments (20 parasites per experiment). Statistical significance was determined by unpaired Student’s *t* test. (C) TgMyoF-mAID parasites ectopically expressing 3xTy-Atg8 under the Atg8 promoter were grown for 12 hours before treatment with ethanol (control) or IAA (TgMyoF KD) for 6h. Cells were stained with anti-Ty (magenta) and anti-Cpn60 antibodies (yellow). Maximum intensity projections of deconvolved images of parasites at interphase, elongation and post-fission show that loss of TgMyoF does not affect Atg8 localization. Scale bar represents 5 μm

### TgMyoF knockdown did not alter TgAtg8 localization with the apicoplast

TgAtg8 is a protein that localizes to the apicoplast outer membrane. Loss of TgAtg8 disrupts the interaction between the centrosomes and apicoplast resulting in asymmetric distribution of the apicoplast to the growing daughter parasites, reminiscent of the phenotype observed after TgMyoF depletion (Fig. 8A-B) [28]. To determine if loss of TgMyoF affected the localization of TgAtg8 at the apicoplast, TgMyoF-mAID parasites transiently expressing 3xTy-TgAtg8 under the endogenous TgAtg8 promoter were treated with ethanol or 500 μM IAA for 6h before fixation and staining with anti-Cpn60 antibody. TgAtg8 localized to the apicoplast at all stages of the apicoplast division cycle and knockdown of TgMyoF did not disrupt TgAtg8 localization (Fig. 8C and Fig. S7) indicating that the loss of interaction between centrosomes and the elongated apicoplast seen in the TgMyoF knockdown parasites is not due to mislocalization of TgAtg8.

## DISCUSSION

The apicoplast is an essential organelle in *T. gondii*. In order to maintain a functional apicoplast, the parasite must execute two distinct and essential cellular processes: Division of this single copy organelle once per cell cycle to ensure each daughter parasite inherits one apicoplast, and trafficking of nuclear encoded proteins to the apicoplast. We investigated the role of actin and an associated myosin motor, TgMyoF, in these processes.

### Loss of MyoF and actin results in the accumulation of TgAPT1 vesicles in the parasite cytosol

Vesicle trafficking is a multi-step process involving vesicle formation and budding from the donor compartment, transport, docking at the target membrane and finally vesicle fusion. While it is recognized that trafficking of nuclear encoded proteins to the apicoplast is a vital cellular process, our understanding of this trafficking process is incomplete and a number of trafficking pathways have been proposed [52]. Using TgAPT1-EmGFP expressing parasites, our data confirms previously published data [14] demonstrating that vesicle formation is coordinated with the cell cycle, with the number of vesicles increasing 7-fold during the elongation phase of apicoplast division. We were not able to definitively determine the origin of these vesicles and we considered two possible hypotheses: First, APT1 vesicles originate at the apicoplast and represent a mechanism for trafficking apicoplast-synthesized lipids to other subcellular locations, or second, these vesicles contain newly synthesized nuclear encoded apicoplast proteins that originate at the ER, and represent a mechanism for trafficking these proteins to the apicoplast as previously proposed [14,48]. A number of lines of evidence led us to favor the latter hypothesis: (1) vesicle accumulation occurs in “apicoplast-less” parasites indicating that vesicle formation is not dependent on the apicoplast itself, as previously observed (Fig. 2) [48]. (2) APT1 protein synthesis and vesicle formation both occur in a short window in the cell cycle when the apicoplast is undergoing division (Fig. 1), (3) recent work has shown that the SNARE proteins TgVAMP4-2 and TgSNAP29 localize to the apicoplast and are essential for the trafficking of nuclear encoded proteins to the apicoplast [53,54], which highlight a role for vesicles in protein trafficking to the apicoplast, and (4) while we cannot exclude the possibility that these vesicles are derived from the apicoplast itself, vesicle formation at and budding from the apicoplast has never been described in the literature. Additionally, chloroplast to ER lipid trafficking in plants occurs by Acyl-coA’s and does not involve vesicle formation [55]. We had hoped our Pulse-Chase assay could be used to determine the origin of the APT1 vesicles and while we did not observe apicoplast derived vesicles (labeled with JF646), we also did not observe Halo-TMR labeled vesicles. We speculate that the pervasive Halo-TMR signal from the ER hampered our ability to visualize APT1-TMR vesicles in the cytosol.

A 12-hour treatment with CD to induce actin depolymerization or a 15-hour treatment with IAA to deplete TgMyoF caused to a significant increase in the number of vesicles in the cytosol at all phases of the cell division cycle (Fig. 2). It is possible that these vesicles are ER-derived but fail to fuse with the apicoplast and thus accumulate in the cytosol during subsequent cell division cycles. Alternatively, loss of these proteins may result in vesiculation of the apicoplast itself.

### MyoF and actin only play a minor role in APT1 vesicle motion

Given the role of MyoF and actin in dense granule transport, we sought to determine if this cytoskeletal system was also required for APT1 vesicle transport. By tracking the motion of individual vesicles in live parasites we demonstrate that APT1 vesicle motion appears predominately diffusive-like. With ∼80% of vesicles moving in a random manner, i.e., those with a straightness value less than 0.5 (Fig. 3). “Directed” movements were defined as those with a straightness value greater than 0.5 and a displacement length greater than 500nm. Only ∼10% of vesicles in control parasites met these criteria. Actin depolymerization or TgMyoF depletion reduced this percentage by ∼50% (Fig. 3). Therefore, we conclude that only a small proportion of vesicles are transported in an actomyosin dependent manner. Given that 90% of vesicles exhibit diffusive motion it is unlikely that the small change in the percentage of vesicles exhibiting directed motor driven movement can account for the large accumulation in vesicles seen after CD and IAA treatment.

### MyoF and Actin control apicoplast dynamics which mediate apicoplast-centrosome association

The sequence of events that occur during the apicoplast division cycle had previously been established via fixed cell immunofluorescence assays or live-cell imaging with low temporal resolution (time between frames on the order of minutes) [27,32] (Fig. 1A). This study reveals new insight into how this sequence of events proceeds (Fig. 9). The first step in the division cycle is the transition from circular to elongated morphology. Elongated apicoplasts with a length <1.5µm were on average ∼1µm away from the centrosomes. Thus, the transition from circular to elongated occurs before, and is not dependent on centrosome association. This transition also occurred in the absence of both actin and TgMyoF, indicating that an as yet unidentified protein(s) must control apicoplast membrane deformation.

**Figure 9.**
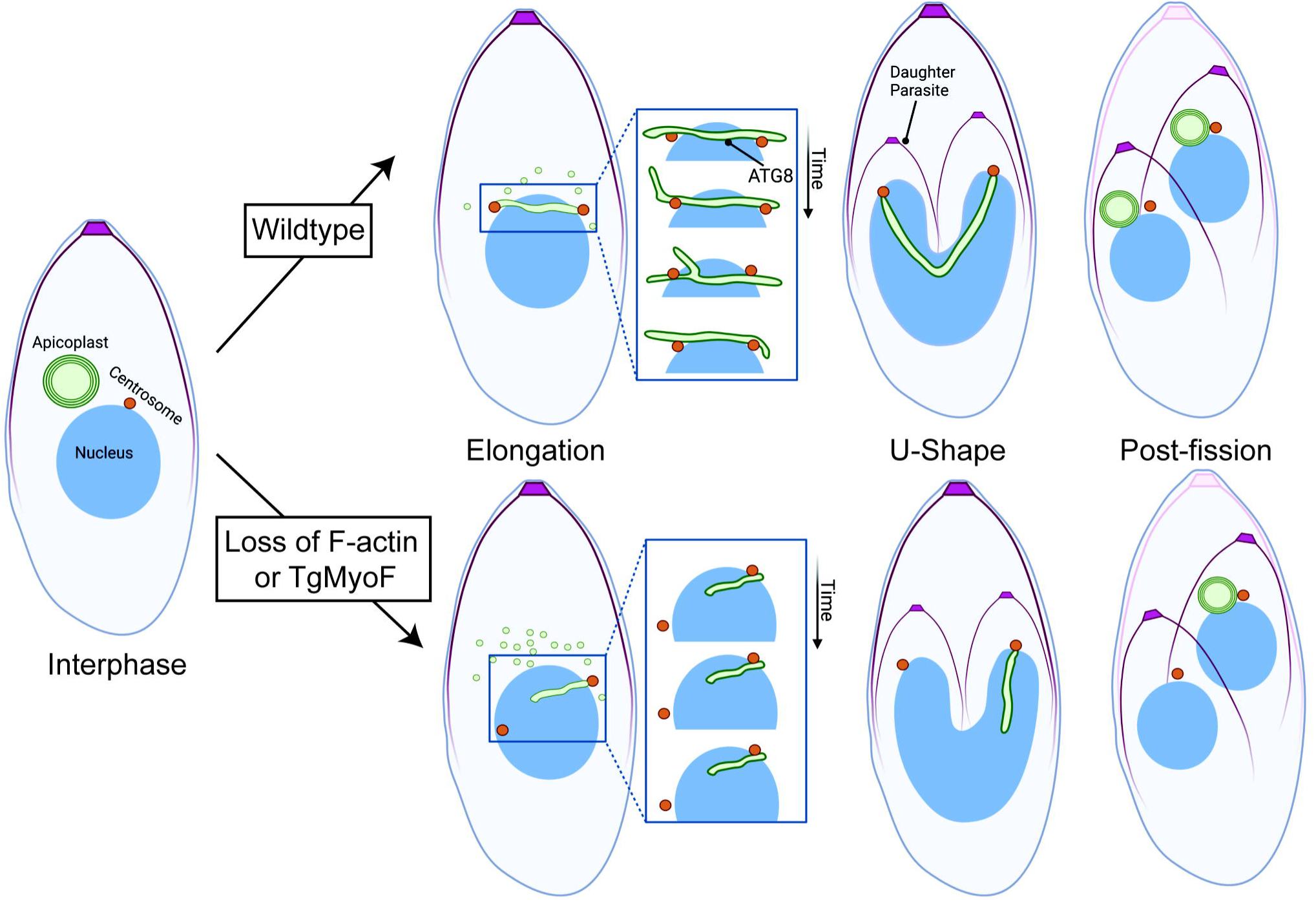
Illustration of the role of actin and TgMyoF in apicoplast division. In interphase parasites, *T. gondii* contains a single apicoplast and centrosome. At the onset of apicoplast division, the apicoplast elongates in a manner independent of centrosome association and there is an increase in the number TgAPT1+ vesicles in the cytosol, which could supply the apicoplast with additional proteins and lipids prior to duplication. In the elongation phase, the apicoplast is highly dynamic exhibiting bending and branch formation. At this stage, the centrosome is dynamically associated with both the sides and the tips of the apicoplast. Association of the apicoplast tips with the centrosome facilitates inheritance of a single apicoplast by each daughter cell upon apicoplast fission. In the absence of TgMyoF and actin, TgAPT1+ vesicles accumulate in the cytosol. These vesicles could represent either vesiculation/fragmentation of the apicoplast or ER-derived vesicles that have failed to fuse with the apicoplast. Also during elongation, the apicoplast is static and the apicoplast frequently fails to associate with one or both centrosomes. This results in the asymmetric division of the apicoplast into daughter parasites.

Since apicoplast elongation is not dependent on centrosome association a mechanism must be in place to drive the interaction of these two organelles. The high-temporal resolution live cell imaging presented here provided insight into how this is achieved. Apicoplast branching, extension and retraction are needed to move the apicoplast in close proximity to the nuclear envelope-embedded centrosomes to facilitate an interaction with either the sides or tips of the apicoplast. Centrosome sliding along the apicoplast side and branch formation and retraction are likely important for positioning of the centrosome at the apicoplast tips. Consistent with this, the localization of TgAtg8 which plays a role in mediating these interactions, is not restricted to the apicoplast tips but found on the entire apicoplast surface (Fig. 8). Loss of TgMyoF results in decreased apicoplast dynamics (Fig. 5) and centrosome association (Fig. 7), which ultimately leads to apicoplast inheritance defects (Fig. 8). Despite the defects in centrosome-apicoplast association in the absence of TgMyoF, TgAtg8 remains localized at the apicoplast. Collectively, this data indicates that actomyosin-driven apicoplast dynamics are important to facilitate the association between these two organelles and for positioning of the centrosomes at the apicoplast tips.

There are still a number of important questions that need to be addressed in future studies. Previous work [35] has demonstrated that the actin nucleator formin 2, relocalizes from the Golgi to the centrosomes during apicoplast division, however it is unclear how F-actin interacts with the apicoplast in order to drive apicoplast dynamics. Since MyoF and actin are not required for apicoplast elongation, another as yet unidentified protein or proteins must drive membrane deformation and the dramatic transition from a circular to elongated morphology that occurs at the start of apicoplast division.

Overall, our results have determined that MyoF and actin control apicoplast dynamics which is needed for apicoplast-centrosome association, a vital step of the apicoplast division cycle.

## ACKNOWLEDGEMENTS

We thank the members of the Heaslip Lab for helpful discussion during the course of these experiments. We thank our colleagues for sharing reagents: Dr. Gary Ward (University of Vermont) for the IMC-1 antibody; Dr. Boris Striepen (University of Pennsylvania) for the anti-Cpn60 antibody. Dr. Ke Hu (Arizona State University) for pTub-mcherry-TubA1 and pTub-eGFP-TubA1, pTub-FNR-RFP plasmids. Dr. Sebastian Besteiro (Univ. Montpellier, France) for the pTub-tdTomato-TgAtg8 expression plasmid. Schematics in figures 1 and 9 were made with Biorender.

## FUNDING INFORMATION

This work was funded by the National Institute of General Medical Sciences 1R35GM138316-01 awarded to ATH. The funders had no role in study design, data collection and interpretation, the decision to submit the work for publication or manuscript preparation. The authors declare that no competing interests exist.

## Notes

### Competing Interest Statement

The authors have declared no competing interest.

### Summary of Updates

Organization of the manuscript has been revised. New data and analysis has been added to Figures 1 and 7.

