## Supplemental Information for "F-actin and Myosin F control apicoplast elongation dynamics which drive apicoplast-centrosome association in *Toxoplasma gondii*"

Devarakonda et al., 2023

### Supplementary Figure Legends

#### Figure S1: Creation of TgAPT1-EmGFP parasite line

(A) Schematic of TgAPT1 genomic locus in parental Ku80 parasites and TgAPT1-EmGFP knock-in parasites. Binding sites of primers used for diagnostic PCR are indicated. Expected PCR product size with F1 and R1 primers is 1785bp. Expected PCR product size for F1 and R2 primers is 1053bp. (B) Diagnostic PCR with F1/R2 and F1/R1 primer pairs using genomic DNA from parental Ku80 and APT1 knock-in parasites as a template.

#### Figure S2: Creation of TgMyoF-mAID::TgAPT1-EmGFP parasite line

(A) Schematic of UPRT genomic locus in parental TgMyoF-mAID parasites and TgMyoF-mAID::TgAPT1-EmGFP knock-in parasites. Binding sites of proteins used for diagnostic PCR are indicated. Expected PCR product size with F1 and R1 primers is 1101bp. Expected PCR product size for F1 and R2 primers is 2373bp. (B) Diagnostic PCR with F1/R1 and F1/R2 primer pairs using genomic DNA from parental and knock-in parasites as a template.

#### Figure S3: Actin depolymerization with cytochalasin D causes an accumulation of TgAPT1 vesicles in the cytosol

Fluorescent images of TgAPT1-EmGFP parasites treated with DMSO (control) or 0.2 $\mu$M cytochalasin D treated for 12 hours that were categorized as interphase, elongation and post-fission stages of the apicoplast division cycle. Parasites were fixed and stained with anti-Cpn60 antibody (a marker for the apicoplast shown in yellow) and DAPI (cyan). Panels are maximum intensity projections of deconvolved images. Scale bar represents

5 μm. White box indicates area used to make inset. Inset: Brightness and contrast of insets was adjusted to facilitate visualization of dimmer TgAPT1 vesicles. Inset scale bar represents 2 μm.

**Figure S4: TgMyoF knockdown causes an accumulation of TgAPT1 vesicles in the cytosol**

Fluorescent images of TgMyoF-mAID; TgAPT1-EmGFP parasites treated with ethanol (control) or 500 μM IAA treated for 15 hours and then categorized as being in the interphase, elongation and post-fission stages of the apicoplast division cycle. Parasites were fixed and stained with anti-Cpn60 antibody (a marker for the apicoplast shown in yellow) and DAPI (cyan). Panels are maximum intensity projections of deconvolved images. Scale bar represents 5 μm. White box indicates area used to make inset. Inset: Brightness and contrast of insets was adjusted to facilitate visualization of dimmer TgAPT1 vesicles. Inset scale bar represents 2 μm.

**Figure S5: TgMyoF knockdown did not lead to changes in apicoplats morphology in interphase of post-fission stages of apicoplast division.**

(A) TgMyoF-mAID parasites expressing eGFP-tubulin (magenta) grown for 12 hours before treatment with ethanol (control) or IAA (TgMyoF KD) for 6 hours. Parasites with an anti-Cpn60 antibody (yellow) and DAPI (cyan). Parasites in interphase and post-fission stages of apicopalst divison are shown. Scale bar represents 5 μm.

**Figure S6: TgMyoF knockdown results in asymmetric localization of elongated apicoplast to daughter parasites.**

Maximum intensity projection of deconvolved fluorescent images of TgMyoF-mAID parasites treated with ethanol or IAA.  Parasites are expressing eGFP-tubulin (magenta) stained with anti-Cpn60 antibody (yellow). Control parasites show that the elongated apicoplasts are in proximity to the daughter parasites (white arrowheads). Later in division, the tips of the elongated apicoplast are localized with the growing daughters (magenta arrowheads). Upon TgMyoF KD, there is an asymmetric localization of the elongated apicoplast to one of the daughter cells (white arrow). Parasites where the apicoplast association with the daughters appears normal is indicates with a white arrowhead. In some parasites the apicoplast is not associated with either daughter (indicated with an asterisk).  Scale bar represents 5 μm.

**Figure S7: TgMyoF knockdown does not disrupt ATG8 association with the apicoplast.**

TgMyoF-mAID parasites ectopically expressing 3xTy-Atg8 under the Atg8 promoter were grown for 12 hours before treatment with ethanol (control) or IAA (TgMyoF KD) for 6 hours. Cell were stained with anti-Ty (magenta) and anti-Cpn60 antibodies (yellow), and DAPI. Maximum intensity projections of deconvolved images of parasites during interphase, elongation and post-fission are shown. Scale bar represents 5 μm

**Table S1: List of plasmids used in this study**

**Table S2: List of primers used in this study**

**Table S3: List of gene accession numbers associated with this study**

**Table S4: List of antibodies used in this study**

**Video S1:** TgAPT1-EmGFP parasites were imaged in the elongation phase of apicoplast division. Images are maximum intensity projection of deconvolved images that contained 3 Z-slices with 0.4µm step size between planes. Imaging speed/stack was 2 stacks/second. Playback speed is 30fps or 15x real time.

**Video S2:** TgAPT1-EmGFP parasites were imaged in the elongation phase of apicoplast division. Examples of apicoplast that are dynamic (left) and static (right) over the imaging period. Images are maximum intensity projection of deconvolved images that contained 3 Z-slices with 0.4µm step size between planes. Imaging speed/stack was 2 stacks/second. Playback speed is 40fps or 20x real time.

**Video S3:** TgAPT1-EmGFP parasites treated with DMSO (left) and CD (right) for 30 minutes were imaged in the elongation phase of apicoplast division. Images are maximum intensity projection of deconvolved images that contained 3 Z-slices with 0.4µm step size between planes. Imaging speed/stack was 2 stacks/second. Playback speed is 30fps or 15x real time.

**Video S4:** TgMyoF-AID: TgAPT1-EmGFP parasites treated with ethanol (left) or IAA (right) for 15 hours were imaged in the elongation phase of apicoplast division. Images are maximum intensity projection of deconvolved images that contained 3 Z-slices with 0.4µm step size between planes. Imaging speed/stack was 0.5 stacks/second. Playback speed is 25fps or 12x real time.

**Video S5:** RH parasites expressing and pmin-Centrin1-eGFP (centrosome marker; magenta) ptub-FNR-RFP (apicoplast marker; green) in the elongation phase of apicoplast division. Images are maximum intensity projection of deconvolved images that contained 5 Z-slices with 0.4um step size between planes. Images were acquired channel 1/channel 2 at each Z slice. Time lag between channels was ~0.4 seconds. Imaging speed is 4.6s/2-color stack. Total imaging time is 9 minutes. Playback speed is 20fps or 84x real time. Raw data was bleach and drift corrected as outlined in materials and methods

**Video S6:** Inset 1 from Video S5/Figure 6. Playback speed is 5fps or 22x real time.

**Video S7:** Inset 2 from Video S5/Figure 6. Playback speed is 10fps or 45x real time.

**Video S8:** Inset 3 from Video S5/Figure 6. Playback speed is 10fps or 22x real time.

**Video S9:** RH parasites expressing and pmin-Centrin1-eGFP (centrosome marker; magenta) ptub-FNR-RFP (apicoplast marker; green) in interphase and post-fission. Images are maximum intensity projections of deconvolved images that contained 5 Z-slices with 0.4μm step size between planes. Images were acquired channel 1/channel 2 at each Z slice. Time lag between channels was ~0.4 seconds. Imaging speed is 7.8s/2-color stack. Total imaging time is 2.5 minutes. Playback speed is 60x real time. Raw data was bleach and drift corrected as outlined in materials and methods

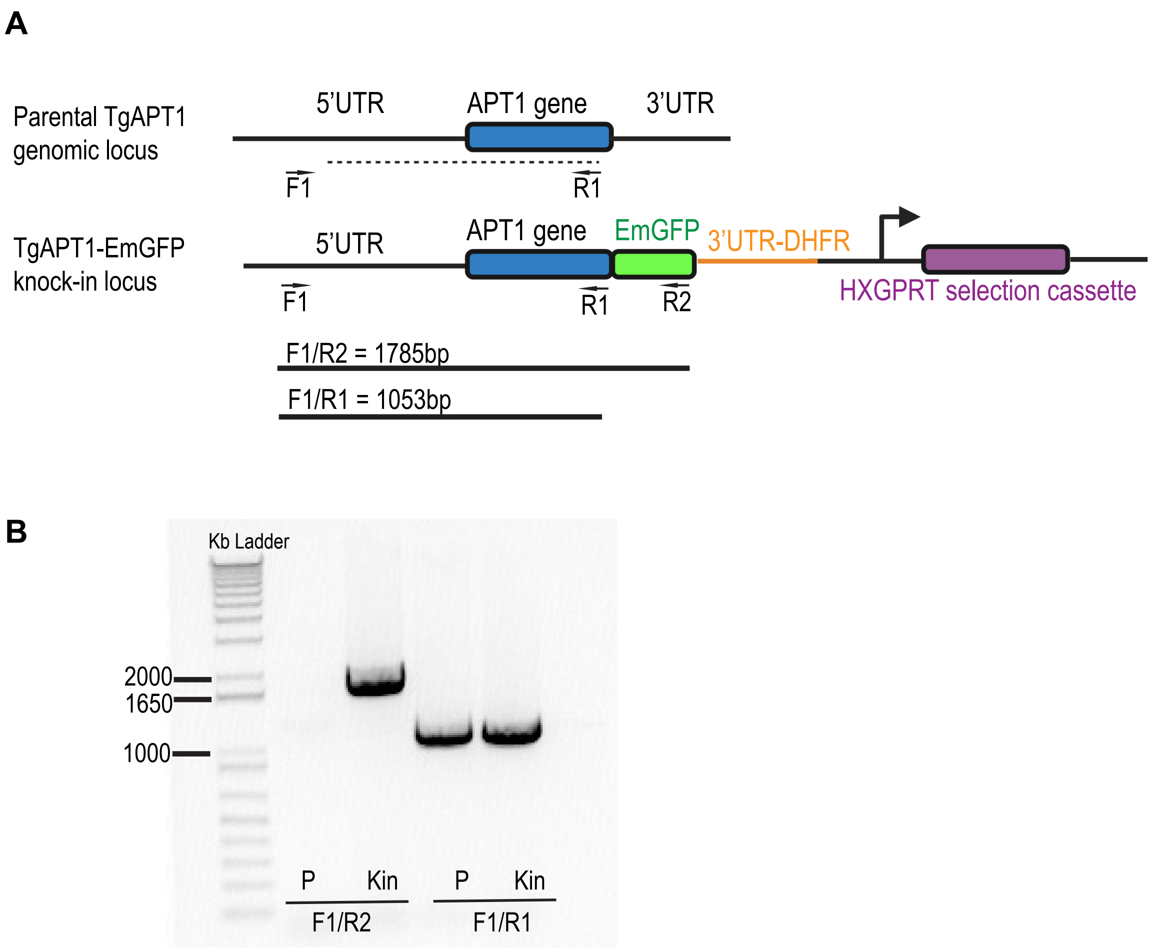


**Figure S1: Creation of TgAPT1-EmGFP parasite line.**

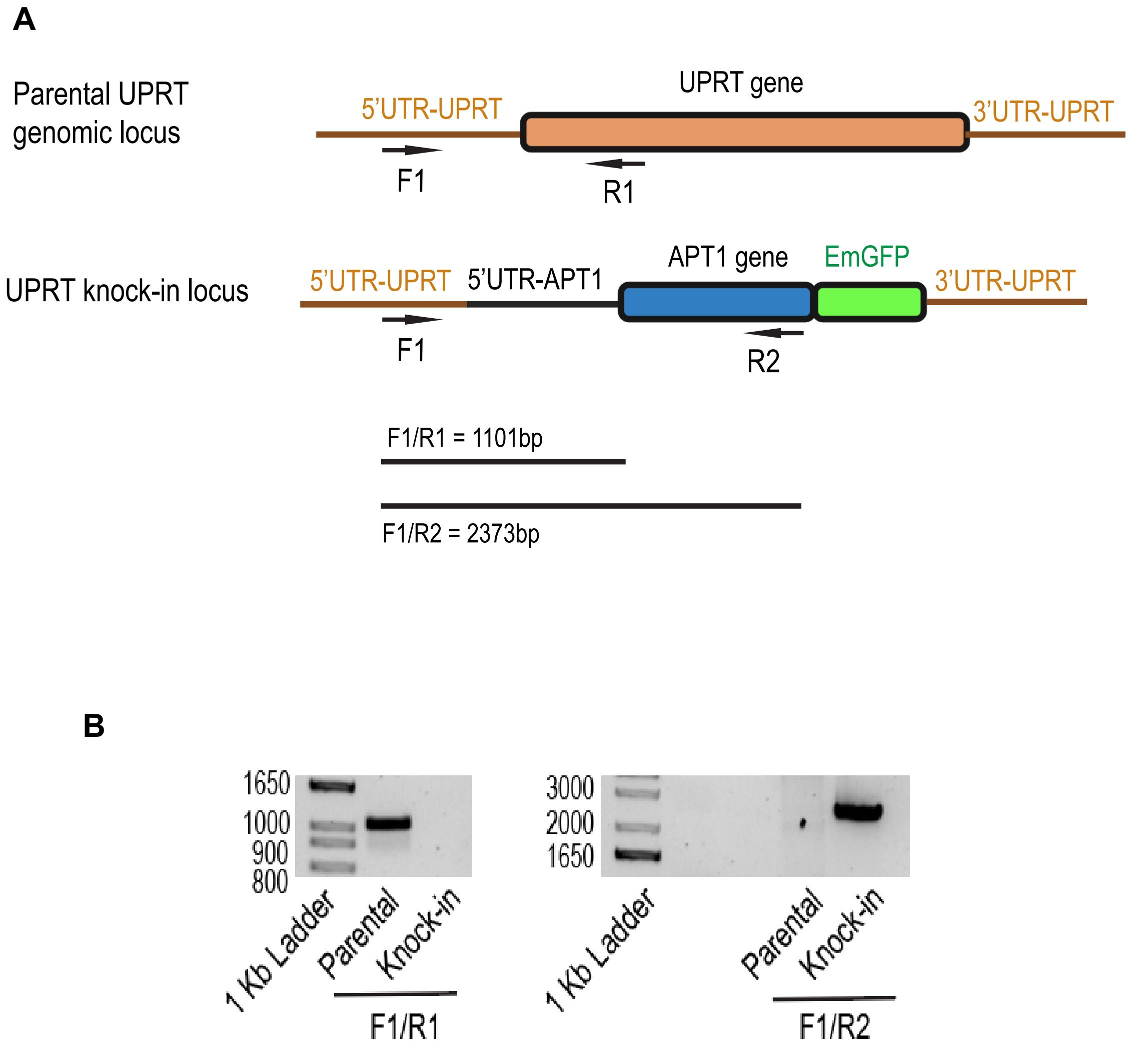


**Figure S2: Creation of TgMyoF-mAID; APT1-EmGFP parasite line**

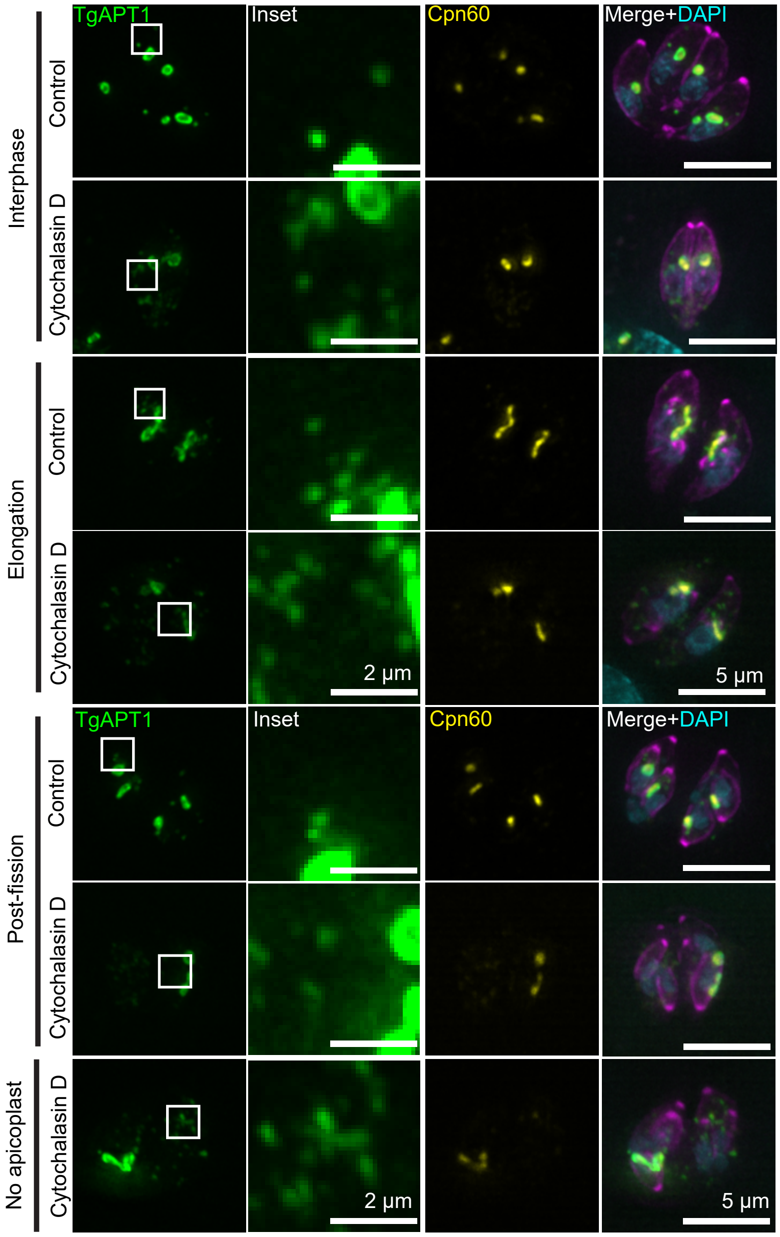


**Figure S3: Actin depolymerization with cytochalasin D causes an accumulation of TgAPT1 vesicles in the cytosol**

Fluorescent images of TgAPT1-EmGFP parasites treated with DMSO (control) or 0.2 mM cytochalasin D treated for 12 hours that were categorized as interphase, elongation and post-fission stages of the apicoplast division cycle. Parasites were fixed and stained with anti-Cpn60 antibody (a marker for the apicoplast shown in yellow) and DAPI (cyan). Panels are maximum intensity projections of deconvolved images. Scale bar represents 5 µm. White box indicates area used to make inset. Inset: Brightness and contrast of insets was adjusted to facilitate visualization of dimmer TgAPT1 vesicles. Inset scale bar represents 2 µm.

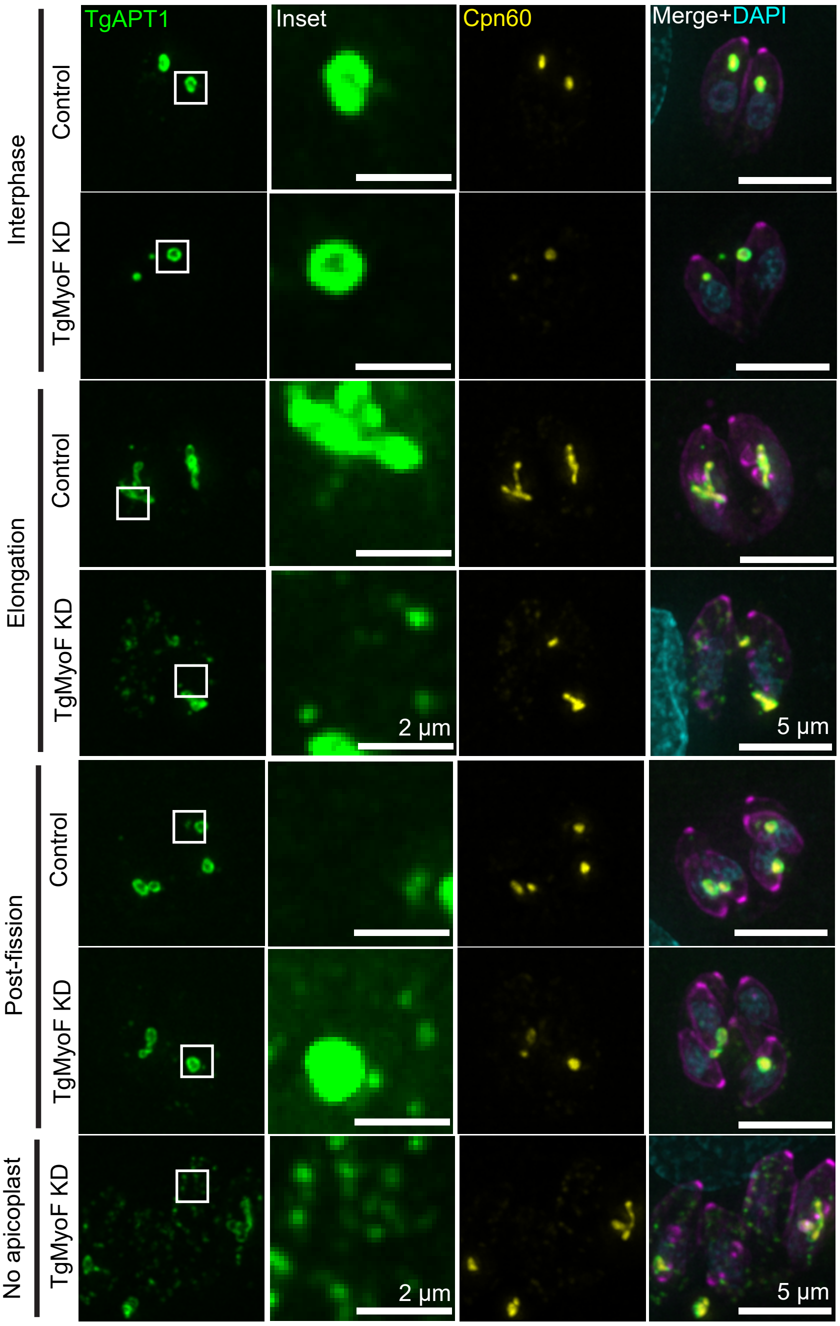


**Figure S4: TgMyoF knockdown causes an accumulation of TgAPT1 vesicles in the cytosol**

Fluorescent images of TgMyoF-mAID; TgAPT1-EmGFP parasites treated with ethanol (control) or 500 M IAA treated for 15 hours and then categorized as being in the inter-phase, elongation and post-fission stages of the apicoplast division cycle. Parasites were fixed and stained with anti-Cpn60 antibody (a marker for the apicoplast shown in yellow) and DAPI (cyan). Panels are maximum intensity projections of deconvolved images. Scale bar represents 5 µm. White box indicates area used to make inset. Inset: Brightness and contrast of insets was adjusted to facilitate visualization of dimmer TgAPT1 vesicles. Inset scale bar represents 2 µm.

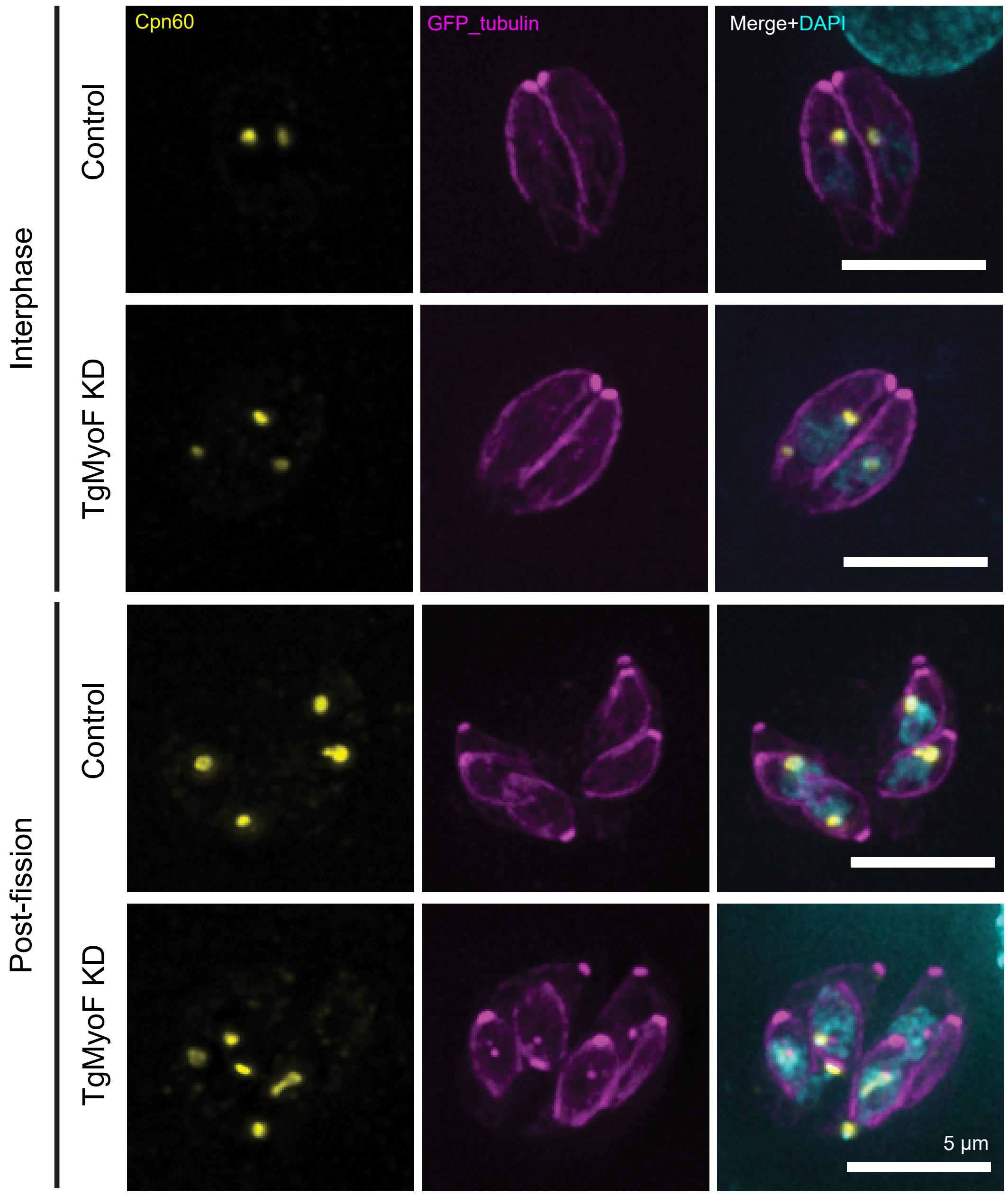


**Figure S5: TgMyoF knockdown did not lead to changes in apicoplats morphology in interphase or post-fission stages of apicoplast division**.
TgMyoF-mAID parasites expressing eGFP-tubulin (magenta) grown for 12 hours before treatment with ethanol (control) or IAA (TgMyoF KD) for 6 hours. Parasites with an anti-Cpn60 antibody (yellow) and DAPI (cyan). Parasites in interphase and post-fission stages of apicopalst divison are shown. Scale bar represents 5 μm.

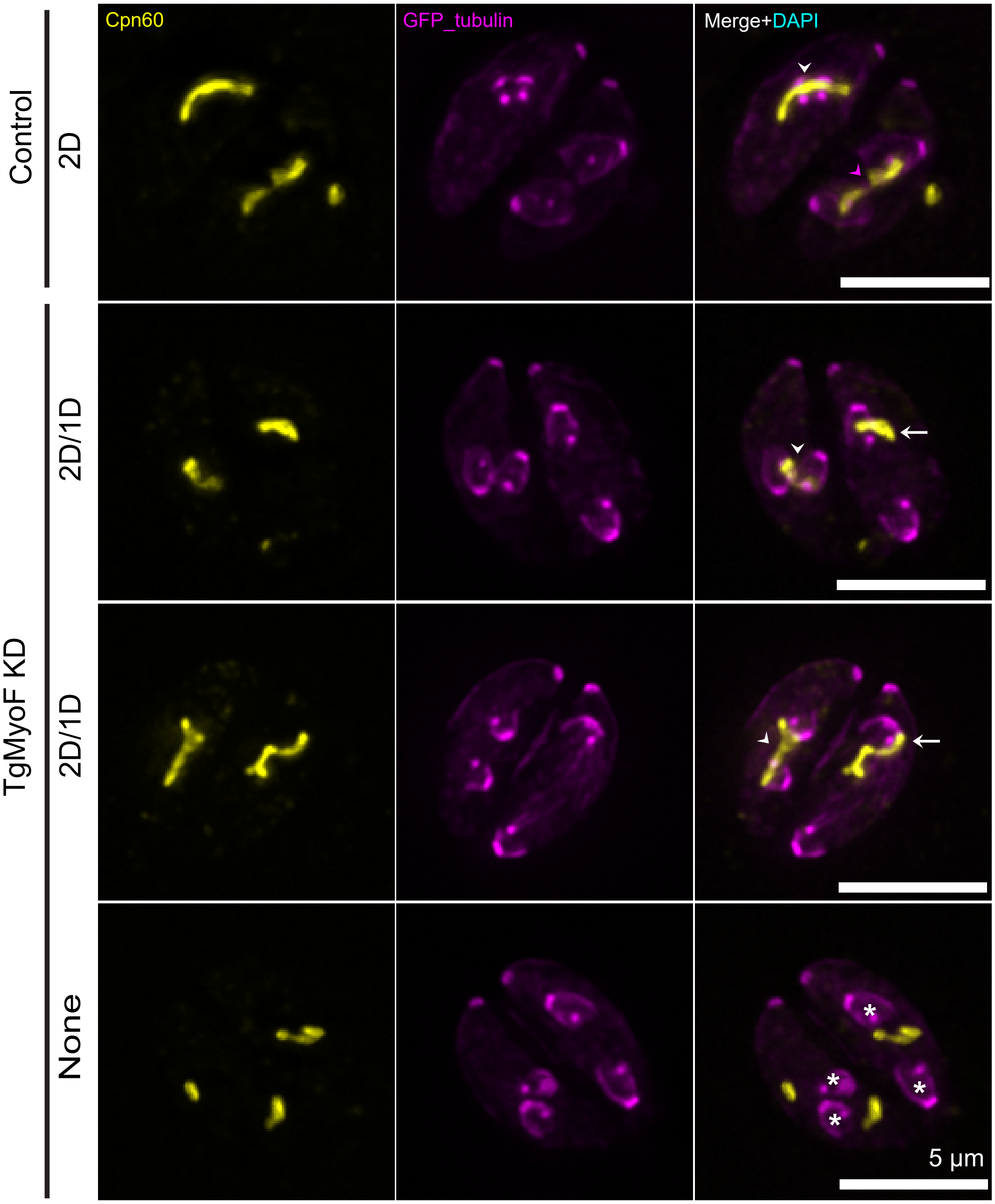


**Figure S6: TgMyoF knockdown results in asymmetric localization of elongated apicoplast to daughter parasites.**

Maximum intensity projection of deconvolved fluorescent images of TgMyoF-mAID parasites treated with ethanol or IAA. Parasites are expressing eGFP-tubulin (magenta) stained with anti-Cpn60 antibody (yellow). Control parasites show that the elongated apicoplasts are in proximity to the daughter parasites (white arrowheads). Later in division, the tips of the elongated apicoplast are localized with the growing daughters (magenta arrowheads). Upon TgMyoF KD, there is an asymmetric localization of the elongated apicoplast to one of the daughter cells (white arrow). Parasites where the apicoplast association with the daughters appears normal is indicates with a white arrowhead. In some parasites the apicoplast is not associated with either daughter (indicated with an asterisk). Scale bar represents 5 µm.

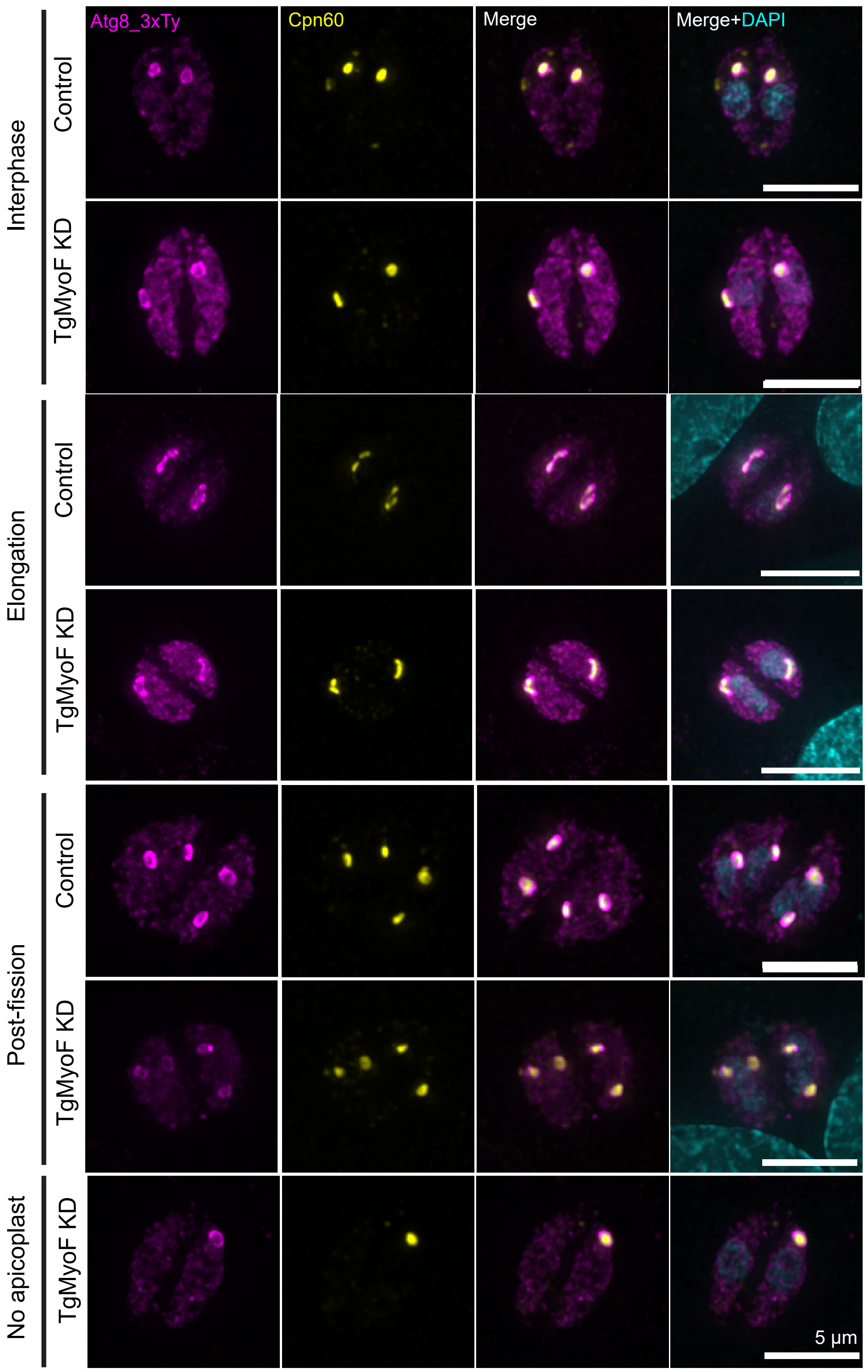


**Figure S7: TgMyoF knockdown does not disrupt ATG8 association with the apicoplast.** TgMyoF-mAID parasites ectopically expressing 3xTy-Atg8 under the control of the Atg8 promoter were grown for 12 hours before treatment with ethanol (control) or IAA (TgMyoF KD) for 6 hours. Cell were stained with anti-Ty (magenta) and anti-Cpn60 antibodies (yellow), and DAPI (cyan). Maximum intensity projections of deconvolved images of parasites during interphase, elongation and post-fission are shown. Scale bar represents 5 μm

**Table 1.** List of Plasmids used in this study

| Plasmid Name | Purpose | Reference |
|---|---|---|
| pTKOII_APT1_EmGFP | Endogenous tagging of APT1EmGFP genomic locus | This study |
| pUPRT_APT1_EmGFP_3'UPRT | Expression of APT1-EmGFP from UPRT locus | This study |
| pUPRT_APT1_Halo_3'UPRT | Expression of APT1-HaloP from UPRT locus | This study |
| pUPRT_pMyoF_mCherry_MyoFCDS_3'UPRT | Expression of mCherry-MyoF from UPRT locus | Unpublished plasmid |
| pAtg8_3xTy_Atg8 | Fluorescent labeling of Atg8 subcloned from ptub_tdtomato_Atg8 (Leveque et al., 2015 PMID: 26507233) | This study |
| pmin_Centrin1_GFP | Fluorescent labeling of centrosomes | Hu K. 2008; PMID: 18208326 |
| ptub_mcherry_ tubulinA1 | Fluorescent labelling of parasite periphery and daughters | Hu et al., 2002; PMID: 11901169 |
| ptub_eGFP_tubulinA1 | Fluorescent labelling of parasite periphery and daughters | Hu et al., 2002; PMID: 11901169 |

**Table S2:** List of primers used in this study

| Primer Name | Sequence | Purpose |
|---|---|---|
| TgAPT1 F1 | AAACGACGGCCAGTGAGCGCGCCCACCG CGGTGGCCTAGGcctcgcgcgcgttcgtccgg | Gibson Primer (F) for inserting APT1 gene into pTKOII//EmGFP Plasmid |
| TgAPT1 R1 | TGAACAGCTCCTCGCCCTTGCTCACGAGT CCGGAAGATCTtccgtacttggtcttcgaga | Gibson Primer (R) for inserting APT1 gene into pTKOII//EmGFP Plasmid |
| TgAPT1 promoter F1 | TACCACTTCGCTTCCCTGTC | PCR Confirmation of integration of APT1-EmGFP into the endogenous locus |
| APT1 CDS R | TCCGTACTTGGTCTTCGAGAGA | PCR Confirmation of integration of APT1-EmGFP into the endogenous locus |
| EmGFP R | TTACTTGTACAGCTCGTCCATG | PCR Confirmation of integration of APT1-EmGFP into the endogenous locus |
| Halo-F1 | AAGACCAAGTACGGAagatctGAAATCG GTACTGGCTTTCCATT | To amplify Halo coding sequence for creating pAPT1-APT1-Halo construct |
| Halo-R1 | aaaaaaactagagaccttaagTTAACCGGAAATC TCCAGAGTAGA | To amplify Halo coding sequence for creating pAPT1-APT1-Halo construct |
| 5'-APT1-UPRT-F | CGTATTCCTTTTTTCGTCGGACCTTTC CACAGGGCCTAGGcctcgcgcgcgttcgtccgg | Gibson Primer (F) for inserting APT1 gene into pTKOII//mcherryMyoF Plasmid |
| 3'-APT1-UPRT-R | ATTCCGTCAGCGGTCTGTCAAAAAAA CTAGAGACCTTAAGttacttgtacagctcgtcca | Gibson Primer (R) for inserting APT1 gene into pTKOII/mcherry MyoF Plasmid |
| 5'-UPRT F | TTTTCCTGTTTTCGTCGTCATC | Creation of CrispR-HR Oligo and PCR Confirmation of integration of APT1-EmGFP into the UPRT locus |
| 3'-UPRT R | TACCACTTCGCTTCCCTGTC | Creation of CrispR-HR Oligo |
| UPRT-Intron R | AGAGAGTTGAGAACAGGCTTC | PCR Confirmation of integration of APT1-EmGFP into the UPRT locus |
| Atg8 promoter F | AAGTGGCGAAACCCTCGGCAGTGCCTTG AAAAGAGGCGCCatgtttttgccctgcggcat | Replacement of pTub with pATG8 to create pATG8-ATG8-tdTomato |
| Atg8 promoter R | TGACCTCCTCGCCCTTGCTCACCCTAGGC ATATGAGATCTgtttttcaagacggcgaata | Replacement of pTub with pATG8 to create pATG8-ATG8-tdTomato |
| 3xTy F | tattcgccgtcttgaaaaacacctaggATGGAGGTCCAT ACTAACCA | Replacement of tdTomato with 3xTy to create pATG8-ATG8-3xTy1 |
| 3xTy R | AGGACACTTCGTCGCGAATCGATGGCAT CCCATGAGATCTatcaagggggtcttggttcg | Replacement of tdTomato with 3xTy to create pATG8-ATG8-3xTy1 |

Table S3: List of gene accession numbers associated for proteins in this study

| **Gene Name** | **ToxoDB Accession Number** |
|---|---|
| TgMyoF | TgME49_278870 |
| TgAPT1 | TgME49_261070 |
| TgATG8 | TgME49_254120 |
| TgCpn60 | TgME49_240600 |
| Centrin 1 | TgME49_247230 |

**Table S4:** List of antibodies used in this study

| Primary Antibody | Catalog # | Dilution | Reference/Source |
| --- | --- | --- | --- |
| Rabbit anti-Cpn60 | Custom | 1:2000 | Dr. Boris Striepen; University of Pennsylvania |
| Mouse anti-IMC1 | Custom | 1:750 | Dr. Gary Ward; University of Vermont |
| Mouse anti-Ty1 | Custom | 1:1000 | Dr. Christopher deGraffienried; Brown University |
| **Secondary Antibody** | **Catalog #** | **Dilution** | **Reference/Source** |
| Goat anti-mouse Alexa Fluor 488 | A11006 | 1:500 | Invitrogen |
| Goat anti-mouse Alexa Fluor 546 | A11030 | 1:1000 | Invitrogen |
| Goat anti-rabbit Alexa Fluor 647 | A21245 | 1:5000 | Invitrogen |
